# Network mechanisms underlying representational drift in area CA1 of hippocampus

**DOI:** 10.1101/2022.11.10.515946

**Authors:** Federico Devalle, Alex Roxin

**Affiliations:** Centre de Recerca Matemàtica, Campus de Bellaterra, Edifici C, 08193 Bellaterra (Barcelona), Spain

## Abstract

Chronic imaging experiments in mice have revealed that the hippocampal code drifts over long time scales. Specifically, the subset of cells which are active on any given session in a familiar environment changes over the course of days and weeks. While some cells transition into or out of the code after a few sessions, others are stable over the entire experiment. Similar representational drift has also been observed in other cortical areas, raising the possibility of a common underlying mechanism, which, however, remains unknown. Here we show, through quantitative fitting of a network model to experimental data, that the statistics of representational drift in CA1 pyramidal cells are consistent with ongoing synaptic turnover in the main excitatory inputs to a neuronal circuit operating in the balanced regime. We find two distinct time-scales of drift: a fast shift in overall excitability with characteristic time-scale of two days, and a slower drift in spatially modulated input on the order of about one month. The observed heterogeneity in single-cell properties, including long-term stability, are explained by variability arising from random changes in the number of active inputs to cells from one session to the next. We furthermore show that these changes are, in turn, consistent with an ongoing process of learning via a Hebbian plasticity rule. We conclude that representational drift is the hallmark of a memory system which continually encodes new information.

---

In area CA1 of the hippocampus a unique pattern of place-cell activity quickly emerges upon exploration of a novel space ^1–5^, and is reliably re-evoked when the animal is returned to a familiar environment ^6–8^. However, the seeming stability of the hippocampal code only holds true on relatively short time scales. Indeed, in familiar environments, both place-cell- and non place-cell activity slowly changes over days and weeks ^9–17^. These long time-scale changes in the neuronal code, dubbed representational drift (RD), have also been seen in other cortical areas, such as parietal, piriform, visual and auditory cortex ^18–22^. The phenomenology of RD is fundamentally similar in all of these cases: the identity of active neurons changes from session to session, although the sparseness of representation is stable. Furthermore, there is considerable heterogeneity in the stability of cells, or how often they take part in the code. It is not known what the mechanism generating the observed drift is, nor what its potential functional role might be.

From a mechanistic perspective, several computational studies have shown that ongoing plasticity can result in changes in neuronal dynamics reminiscent of RD at the population level ^23–25^. In these studies the plasticity acted as a source of noise, driving changes in the representation of already stored patterns. In fact, it was previously hypothesized that such changes might provide the substrate for a time-stamp of a given memory ^10^. Alternatively, it has also been hypothesized that RD may occur due to plasticity related to the encoding of new memories ^19^. In this case RD would be the unavoidable signature of ongoing learning ^26–28^. However, it remains unclear to what extent these, or other mechanisms can provide a quantitative description for the statistics of RD which are observed experimentally. We therefore sought to develop a biologically plausible computational model which could be fit to data quantitatively, and which specifically allowed us to address questions regarding the underlying mechanism, as well as the role of RD.

Here we elucidate a network mechanism which can account for the observed RD in area CA1 of the hippocampus in mice. We do this by making quantitative fits of a large-scale network of spiking neurons to data from chronic Ca2^+^-imaging experiments ^10^. We infer that changes in neuronal activity from session-to-session are inherited from changes in the afferent inputs to cells in CA1 from their two main sources of excitatory drive, CA3 and layer III of Entorhinal Cortex (EC). In the network model these changes can be due to an increase or a decrease in the number of synaptic inputs, or changes in the presynaptic patterns of activity, or a combination of the two. Interestingly, these changes are entirely random and uncorrelated from cell-to-cell, and follow simple Gaussian statistics. The time-scales of these changes, which we will refer to as plasticity, are distinct along the two pathways: the characteristic timescale of the evolution of EC inputs is two days, while that of CA3 inputs is on the order of a month.

How are the plasticity mechanisms we identify for CA1 relevant for RD more broadly? Because of differences in anatomy and circuit function, we should expect the details of any such mechanism to vary across cortical areas. However, we also should not miss the forest for the trees. Although the physiological details may differ, it may be that the computational mechanism itself is the same. In this vein we show that the plasticity estimated from the fit to the data in CA1 is consistent with the ongoing encoding of patterns of activity via a Hebbian plasticity rule. In short, RD may reflect the interference of newly encoded memories with older ones, a phenomenon long-predicted theoretically ^26,27^. Our work therefore suggests that representational drift should be a ubiquitous phenomenon in cortical areas involved in the continuous encoding of information, such as the formation of episodic memories.

## Results

In order to identify network mechanisms underlying RD in CA1, we first sought to characterize how spatially tuned versus untuned features of neuronal activity differentially contributed to the observed drift. Drift in the population activity in CA1 place cells can be visualized by ordering the activity of the cells by place-field position on a given session ^9^. Using this same ordering for subsequent, or previous, sessions reveals RD if there is a change in spatial tuning or in the overall level of activity of a cell, e.g. if a cell becomes silent. In the case of mice running on a linear track, using the ordering from the first session shows that there is significant drift on subsequent sessions, Fig.1a. In particular, the drift observed by comparing all even trials on session 1 with session 2 (panels 1a and 2) is visibly larger than the within-session variability comparing even and odd trials in session 1 (panels 1a and 1b). We asked to what extent this drift was due to shifts in the place-field location of cells or simply to changes in rate, Fig.1b. To do so we normalized the time-averaged spatial profile of each place cell by its maximal firing rate on each session. Doing so revealed much less drift, and specifically there was no longer a large difference between the drift from session 1 to session 2 and the degree of variability within session 1, Fig.1c. To quantify this effect we compared scatter plots of the mean event rate of all cells within session 1, and then between sessions 1 and 2, Fig.1d. The correlation of the mean rates evaluated on only even trials versus only odd trials within the session 1 was much higher than the correlation of mean rates between sessions 1 and 2. The pronounced drop in correlation occurred reliably from one session to the next regardless of the session identity, as can be seen in the matrix of rate correlations in Fig.1e. The correlation as a function of session separation averaged over all sessions showed a steep initial drop, followed by a slower decay, Fig.1f. In fact, the curve was much better fit by a double exponential than a single exponential suggesting the presence of two processes operating at distinct time-scales.

**Fig. 1.**
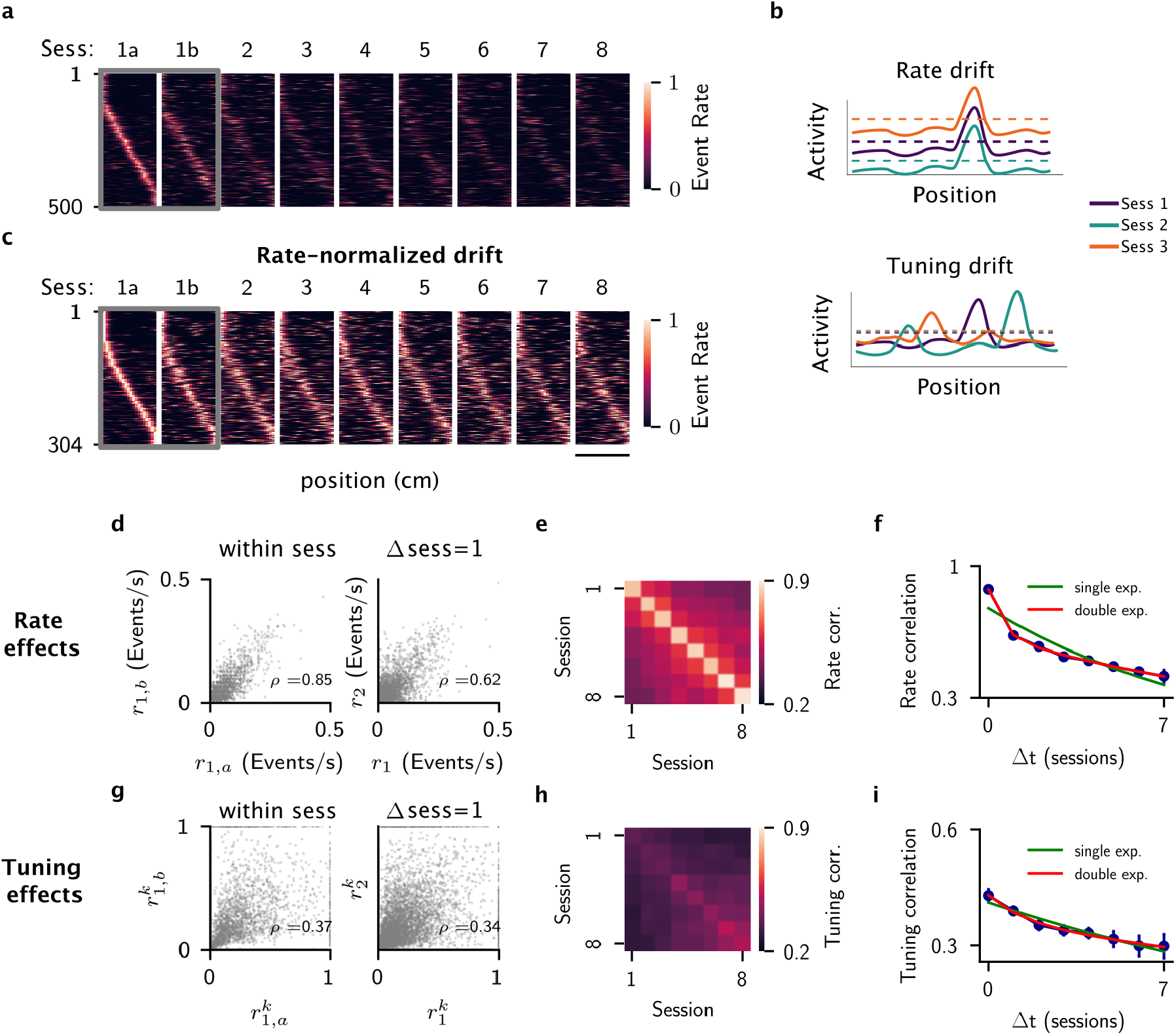
Changes in mean rate and spatial tuning differentially affect RD in area CA1 of hippocampus. **a**, Place field maps of cells active on even trials on session 1. 1a and 1b stand for even and odd trials of session 1. Cells are ordered according to their peak rate position in session 1a. Each cell event rate is normalized to the maximum reached over the 8 sessions. The scale bar is 80cm. **b**, Illustration of different changes Ca1 cells may undergo over time. In the top panel, only the mean rate changes from session to session, while tuning is preserved. In the bottom panel, mean rate is preserved while tuning changes over time. **c**, Same as **a**, but plotting cells active on all sessions, and normalizing the event rate within each session. **d**, Scatter plots of mean event rates for all cells. Left: mean event rate on odd trials (ordinate axis) versus mean event rate on even trials. Right: mean event rate on session 2 (ordinate axis) versus mean event rate on session 1. *ρ* is the Pearson correlation coefficient. **e**, Color map showing the rate correlation for all possible session pairs. The bright diagonal indicates strong within-session correlation compared to across-session correlation. **f**, Mean rate correlation as a function of elapsed sessions (blue points), averaging over all possible session pairs. A double exponential fit (red) reveals the presence of two time scales in the data. **g** Scatter plots of position-specific event rates. Left: event rate in bin *k* on odd trials (ordinate axis) versus the event rate in the same bin on even trials. Right: event rate in bin *k* on session 2 (ordinate axis) versus the event rate in the same bin on session 1. **h**, Same as **e**, but for tuning correlation. **i**, Same as **f**, but for tuning correlation. Note that a single time scale captures the decay over time of the correlation.

On the other hand, when we compared mean rates for each spatial bin of the rate-normalized profiles, we found a much smaller decrease in correlation, Fig.1g. Again this drop was slight from one session to the next irrespective of the session identity, and the curve of the average correlation as a function of session separation showed a shallow decay which was much better fit by a single exponential function than the non-normalized curve, Fig.1h-i. The time-scale of this decay was the same as that of the slower of the two time-scales of the mean rates in Fig.1f. This strongly suggested that the slow time-scale decay in the overall RD was due to changes in the spatial tuning of cells, while the fast drop-off in correlation from one session to the next was related to changes in mean activity. Note, however, that some of this drop in correlation from one session to the next may be attributable to recording instabilities from removal and reattachment of the microscope.

### Statistical Model

We hypothesized that these two time-scales might reflect plasticity along the two main afferent excitatory pathways to CA1: Schaffer collaterals from CA3, and inputs originating from layer III of entorhinal cortex (EC). To test this we investigated if changes in synaptic input alone could account for key experimental findings in a simple, statistical model. Specifically, we considered a population of binary neurons, each one of which received two inputs on any given session *t*, which we call *x*^*t*^ and *y*^*t*^, Fig.2a. On the first session we distributed these inputs randomly across neurons according to Gaussian distributions with zero mean and variances 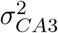 and 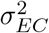 respectively. This led naturally to a large degree of heterogeneity in the inputs to single cells, with some receiving strong excitatory drive and others being strongly inhibited. The total input to a cell *z*^*t*^ = *x*^*t*^ + *y*^*t*^ was then compared to a threshold *θ* to determine if the cell was active or not, i.e. cell activity was binary. Therefore, in this simplified model we did not take continuous changes in the event rate of the cells into account, nor did we explicitly include spatial tuning. To model RD we drew these inputs anew from session to session from the same Gaussian distributions, but allowed for some temporal correlation. Namely, the session-to-session autocorrelation for inputs from CA3 was

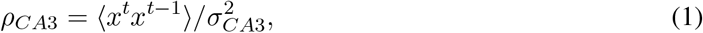

**Fig. 2.**
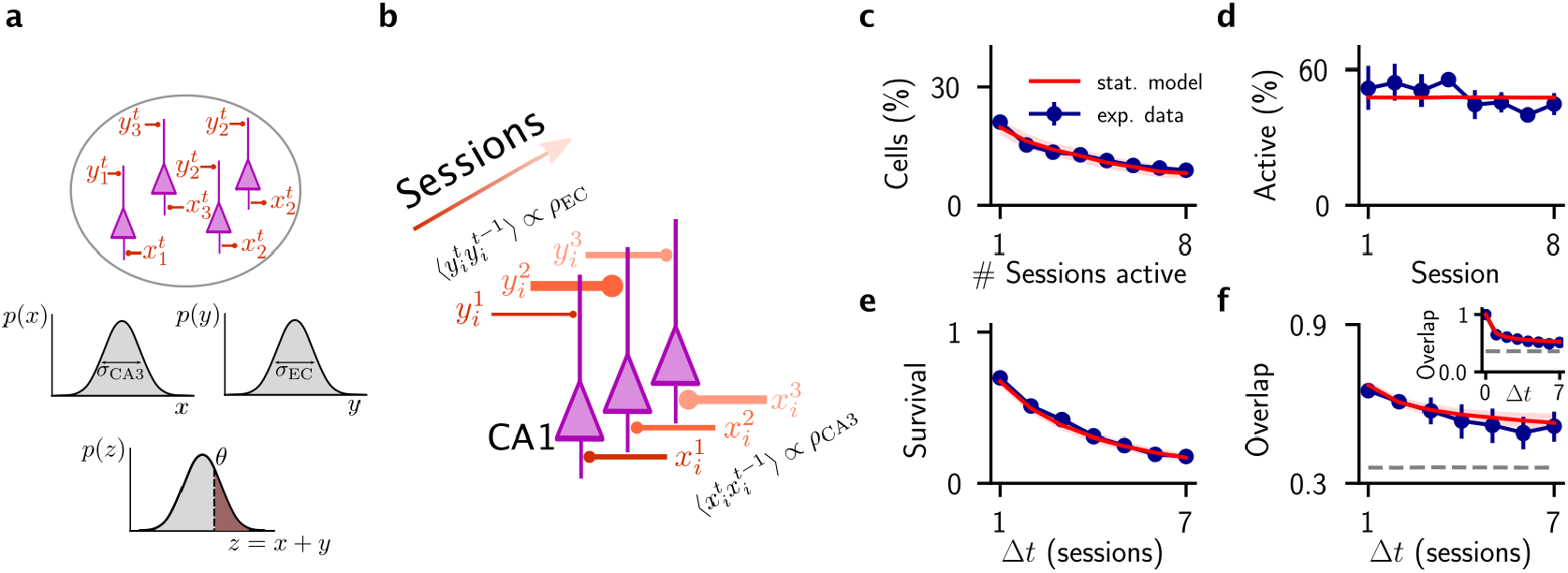
Modeling session-to-session changes in two sources of input with disparate temporal correlations accounts for data on active cells. **a**, A pool of CA1 cells (in pink), where each cell *i* receives two distinct inputs 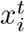 and 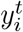 in each session *t*. Inputs are Gaussian distributed across cells with variances 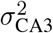 and 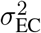 respectively. Cells are active whenever their total input 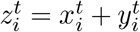 exceeds a threshold *θ*. **b**,The inputs change from session to session with autocorrelations *ρ*_CA3_ and *ρ*_EC_ respectively, while preserving overall input statistics (see Methods). **c-f**, Best fit of the model (red line) to the experimental data (blue points): **c**, Distribution of the number of sessions each cell is active; **d**, Fraction of active cells over time; **e**, Survival probability; **f**, Overlap of the population activity vector. Parameters: *θ* = 0.55, *σ*_*CA*3_*/σ*_*EC*_ = 1.16, *ρ*_CA3_ = 0.95, *ρ*_EC_ = 0.35.

where the brackets indicate an average over all neurons, Fig.2b, and there is an equivalent relation for inputs from EC. There were four model parameters in total after normalization: the ratio of the variances, the threshold, and the two autocorrelations. We optimized these parameters to fit four characteristic measures of neuronal activity from the experimental data: 1 - the distribution of the number of sessions in which a cell is active, 2 - the fraction of active neurons per session, 3 - Survival fraction. This measure includes only those cells which are initially active in the first session, and quantifies the fraction of these which continue to be active on subsequent sessions, and 4 - the overlap in the vector of active neurons from session to session. The model was able to reproduce these measures quantitatively Fig.2c-f, but only when the temporal correlation along one input pathway was close to one (*ρ* = 0.95) and the other sufficiently small (*ρ* = 0.35), Extended Fig.2. In this regime, one of the two inputs changed only very slowly from session to session while the other was almost entirely redrawn randomly for each session, consistent with the two disparate time-scales we observed from direct analysis of the data. Because the model only considered mean inputs, and cell activity was just the sum of the two, it was not possible to say which of the two inputs should vary slowly over time, and which quickly. Once spatial tuning was taken into account (see next section), fast plasticity in EC inputs and slow plasticity in CA3 inputs was the only choice compatible with the data. Finally, we note that a small fraction of cells in the data were active on all 8 sessions, Fig.2c, raising the possibility of a sub-population of stable cells which reliably participated in the hippocampal code. We modeled this by assuming that a fraction *f*_*s*_ of cells were always active, and then allowing the remaining fraction 1 − *f*_*s*_ to evolve according to the model described above. We found that the mean squared error of the model fit simultaneously to the four above-mentioned measures increased for increasing *f*_*s*_, but that it remained low when the stable fraction was below 10 percent, Extended Fig.3. Therefore, while the model fit was always best without dedicated stable cells, the data would nonetheless also be consistent with a small fraction of stable cells.

### A spiking network model reproduces drift statistics in CA1

Having established how the two time-scales of drift could arise via changes in synaptic input quantitatively through a simple statistical model of binary neurons, we next sought to model RD with a more biophysically plausible network model. This allowed us not only to model changes in the firing rate (and hence event rate at the level of Ca2^+^-imaging) of cells as well as their spatial tuning, but also to pinpoint the specific network mechanisms leading to the drift and make predictions which could be tested through additional data analysis. We modeled a population of pyramidal cells and inhibitory interneurons in area CA1 with leaky integrate-and-fire neurons. These CA1 neurons received excitatory input from cells in CA3 and layer III of EC, which were modeled as Poisson processes, Fig.3. We assumed that a fraction *f*_*s*_ of cells in CA3 were spatially tuned in any given environment, which for simplicity we took to have a ring topology. Each cell in CA1 received sparse, random connectivity from a large population of CA3 cells, such that the average number of inputs *K* was large but, in general, much smaller than the total number of presynaptic cells *N*. The strength of synaptic weights was scaled so that the total excitatory input from CA3 was much larger than the distance to threshold for spiking, see Methods for details. Inputs from EC were also random, sparse, and strong in the same sense, and EC cell activity was assumed to be spatially untuned. Although many entorhinal cells exhibit spatial tuning, EC inputs to CA1 pyramidal cells have been found to be largely weakly spatially tuned ^29,30^. Inactivation of Schaffer collateral inputs via optogenetic stimulation largely eliminated place fields in CA1, suggesting a majority contribution of spatial tuning from CA3^31^, although see ^32^. In the parameter regime for which the network model eventually fit the experimental data, adding moderate spatial tuning to EC cells did not significantly affect the results, see Extended data Fig.6. Connections from CA1 pyramidal cells to inhibitory interneurons and vice-versa were also sparse, random, and strong and there were no recurrent excitatory connections, see Methods for details of the network topology.

**Fig. 3.**
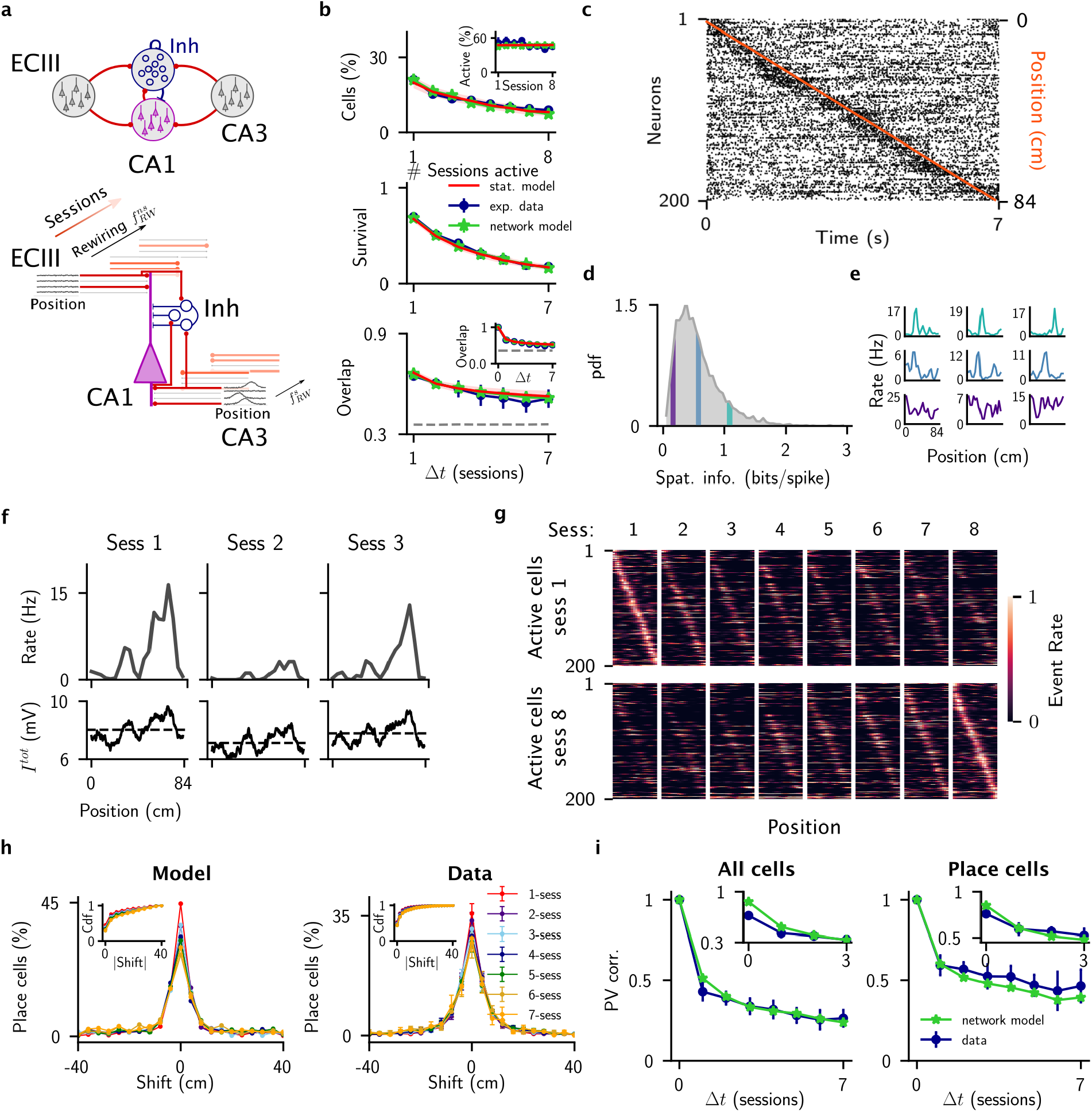
A spiking network model of representational drift in the hippocampus. **a**, Model architecture. Top: connectivity structure of the model. Bottom: illustration of the changes over time of the inputs to a single CA1 cell. **b**, Fit of the network model to the experimental data. **c**, Raster plot of 200 randomly selected cells as a virtual animal runs one lap along the track. In orange the position of the animal superimposed to the spikes of the cells (black dots). **d**, Distribution of spatial information of CA1 exc cells over one session. **e**, Examples tuning curves for 9 selected cells. Color-coded is the value of the spatial information of the example cells. **f**, Tuning curve (top row) and total input (bottom row) for one example cell over three sessions. The dashed line in the bottom panel indicates the average total input along the track for each session. **g**, Place field maps for 200 randomly selected active cells found on session 1 (top) or session 8 (bottom), ordered according to their place field positions. **h**, Distribution of the centroid shift for different number of elapsed sessions (color-coded). Inset: cumulative distribution of the absolute shift. **i**, PV correlation of all cells (left), and only place cells (cells significantly spatially tuned in both sessions) (right). The insets show the first four points of the respective curves, where the initial point (within-session correlation) is computed considering odd vs even trials.

The CA1 network operated in a balanced regime in which the large excitatory and inhibitory input currents to cells cancelled in the mean, leaving the average membrane potential of most cells below the threshold for spiking. Spikes were therefore generated by fluctuations in the membrane potential, resulting in irregular spiking activity and broad, lognormal-like firing rate distributions Extended Data Fig.4, as observed in-vivo ^33,34^. By virtue of the quenched variability in the number of connections per cell, the mean inputs from CA3 and EC were both approximately Gaussian distributed across the network. We fixed the number of inhibitory inputs to each CA1 pyramidal cell so that the mean inhibitory drive to cells was the same and served only to cancel out the mean excitatory input without adding to the variability across cells. Doing this allowed us to map the network dynamics onto the simpler statistical model for a given session straightforwardly since we needed only to determine the variances in the two inputs, which could be calculated analytically, see Methods. In order to model changes in inputs from session to session we explicitly allowed for plasticity in the afferent excitatory synapses from CA3 and EC as well as changes in the patterns of presynaptic activity. Specifically, we find that the autocorrelation of inputs from area CA3 can be written

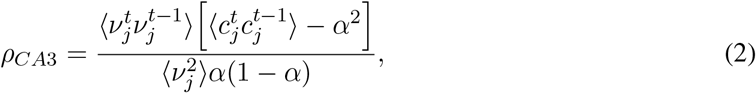

where *ν*_*j*_ is the mean rate of CA3 neuron *j, c*_*j*_ ∈ {0, 1} indicates the absence or presence of a synaptic connection from cell *j, α* is the connection probability, and the brackets indicate an average over the population of CA3 cells. A change in the pattern of presynaptic activity would imply that 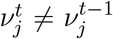 for some neurons *j*, while plasticity implies that 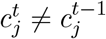 for some neurons *j*. There is an equivalent equation for the autocorrelation of inputs from EC, see Methods for details. Therefore, we were able to match the autocorrelations of inputs in the network model, e.g. Eq.2 to those from the statistical model, e.g. Eq.1 by allowing for changes in connectivity and/or presynaptic activity. In our simulations here we did this by randomly rewiring a fraction of the afferent inputs, while holding the presynaptic activity patterns fixed. In this way we set the network dynamics at a working point close to the optimal fit of the statistical model to the experimental data and with slight parameter adjustments were able to fit the network model as well, Fig.3b.

The network not only reproduced the statistics related to active cells, it also generated place-cell activity in CA1 cells which we could compare to additional experimental data. We modeled the motion of a virtual animal on a circular track by driving spatially selective cells in CA3 with an external signal which represented the animal’s position, see Methods for simulation details. Cells in CA1, which received a large number of random inputs from CA3, also exhibited spatially modulated activity, which closely tracked the true position of the animal Fig.3c. The distribution of spatial information of CA1 cells was broad, with some cells exhibiting sharply peaked place fields, while a majority had much more weakly-tuned profiles Fig.3d-e. The emergence of place-field activity in CA1 occurred as a consequence of the balanced state of the network; the large untuned component of the total excitatory and inhibitory inputs canceled in the mean, leaving a subthreshold spatial tuning on the order of the distance to threshold, which drove spiking. A similar mechanism has been invoked to explain the emergence of orientation selectivity in primary visual cortex in the absence of hypercolumns ^35^.

We asked how the rewiring of inputs from CA3 and EC affected the patterns of activity in the network during exploration of the same environment on separate sessions. Here we first consider that the EC inputs evolve on the faster time-scale, while the CA3 inputs change much more slowly. Typically, significant rewiring of spatially untuned inputs from EC from one session to the next resulted in a net shift in the DC depolarization of place cells in CA1, Fig.3f. This led to an iceberg effect in which the location of maximal activity was relatively unaffected, but the width of the place field and overall firing rate were strongly modulated. This effect was responsible for the sharp decrease in correlation of the population activity between adjacent sessions in our network model, Fig.3g, and resembled the experimentally observed decorrelation, Fig.1a. The fraction of rewired CA3 inputs, of which a fraction *f*_*s*_ were strongly tuned, was significantly less, and led to a slow drift of the place field locations. This was responsible for the slow decay in correlation of the population activity over sessions seen in Fig.3g. A histogram of the resulting shift in the centroid of place cells over sessions in the network is qualitatively similar to the experimental data Fig.3h, although the overall fraction of place cells in the model was always greater than that seen in experiment. Note that if we assume that it is the CA3 inputs which evolve on the fast time-scale, then the resulting histograms of the centroid shift exhibit a rapid broadening (not shown) which is not compatible with experimental observations. Finally, we considered the correlation of the population vector (PV), which in contrast to the data in Fig.3b included the event rate and spatial profile of cell activity. To compare the firing rate of cells in the spiking network with the event rate estimated from the fluorescence of the Ca2^+^-signal we smoothed the spiking activity with an exponential kernel and detected calcium events whenever the smoothed signal crossed a threshold, see Methods for details. Overall, the network provided a good quantitative fit to the data, both including all cells, as well as for only those cells with significant spatial tuning Fig.3i. We found a significant discrepancy between the model and experiment when considering the correlation of the PV within a session by comparing the one half of trials with the other half, e.g. odd versus even trials. In this case the variability in experiment was always larger, see inset in Fig.3i. This may be attributable to the highly non-stationary activity of CA1 cells in-vivo ^36^, which we did not explicitly model.

### Experimental data shows signatures of hypothesized network mechanisms

In the network model, RD was caused by changes in the excitatory drive to cells, which were random and uncorrelated with the level of activity of the cells on any given session. Specifically, the probability of a given increase or decrease in total synaptic input from one session to the next did not depend on the firing rate of a cell on that session. Given this, the probability that a cell became silent and vanished from the code was always higher if its firing rate was already initially low. Comparing a high-rate cell with a low-rate cell in the model showed that the former were more stable in the face of ongoing changes in input Fig.4a. We quantified this effect in the network model by calculating the probability that a cell was active on session *t* as a function of its firing rate on session *t* − 1 and found a sigmoid-like relationship Fig.4b (left). We found a similar trend in the experimental data Fig.4b (right). There was also a positive correlation between the mean firing rate of a cell (calculated only for active sessions) and the number of sessions in which that cell was active. In the case of the model this correlation increased more or less linearly, while in the data it only showed an increase for cells which were active on at least half of the sessions Fig.4c. One possible confounding factor in making this comparison between the model and experiment is the fact that underlying firing rates in CA1 cells are not steady over single sessions (or even trials), as in the model, but rather vary in time ^36^. Such non-stationarities in cell activity are likely due to modulatory inputs related to contextual cues, and not to plasticity-related changes in inputs. Therefore the mean firing rate, averaged over the session, should still reflect the total mean input, which is the factor driving changes in rates in Fig.4b. Additionally, noise in the recording is expected to bias detection for active cells to those with higher firing rates, which might partially account for the trend in Fig.4b.

**Fig. 4.**
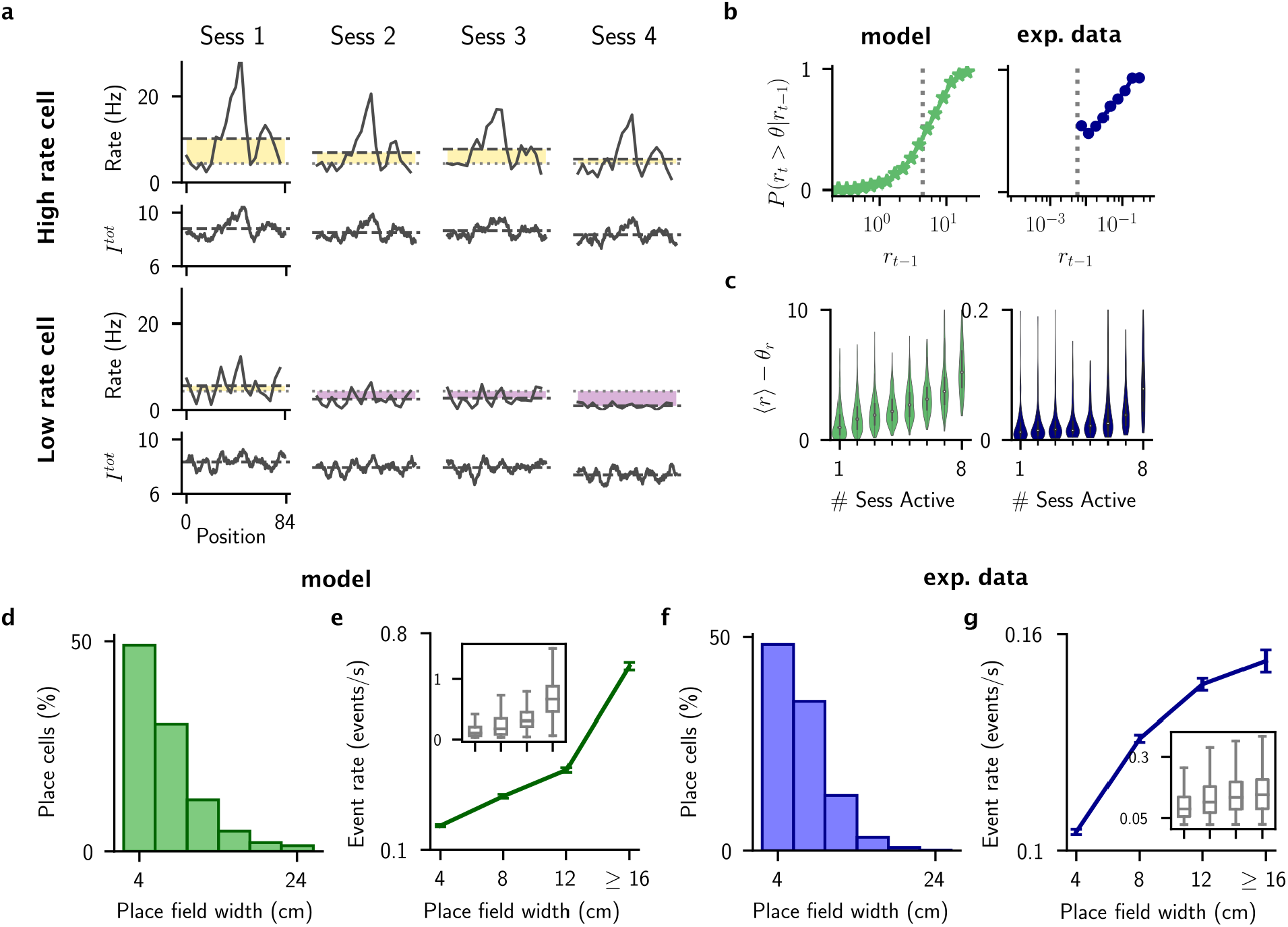
Mean firing rate of cells predicts their stability. **a**, Firing rate and mean total input for two examples cells with high (top) and low (bottom) mean firing rate. Dashed lines indicate averages over all spatial positions, while the light gray dotted line is the threshold. Yellow and purple shadings highlight the distance from threshold, from above and below respectively. **b**, The probability a cell is above threshold given its mean firing rate on the previous session, in the network model (left) and experimental data (right). **c**, Distributions of mean firing rates across sessions, for cells active different number of sessions, in the network model (left), and experimental data (right). **d-g** Statistics of place-field widths in the model and in experiment. **d** Histogram of place field widths in the network model. **e**, Mean event rate of cells grouped by their place field widths in the model. Mean ± s.e.m. Inset shows boxplots. **f-g** Same as **d-e** for the experimental data.

This same network mechanism for RD predicted that place cells should undergo a broadening of their place fields as their mean firing rate increased, as can be seen illustratively in Fig.4a (top). This was due to an iceberg effect for which an increase in net depolarization caused by plasticity would push a tuned subthreshold profile closer to threshold, allowing for spiking activity in the flanks of the former place field. Similarly, decreases in net inputs would silence the flanks, leading to a shrinking of the place field width. We looked for this effect in the network and data. In both cases we found a broad distribution of place-field widths Fig.4d-e, as well as the predicted positive correlation between the activity of the place cells and the width of their place field Fig.4f-g.

### Representational drift is consistent with ongoing Hebbian plasticity

We fit the experimentally observed RD in populations of CA1 cells by randomly rewiring a fraction of inputs to cells in the network model from session to session. We hypothesized that these changes in input connectivity to CA1 could be consistent with plasticity underlying the ongoing encoding of new information. If this were true, then RD would represent the interference of new patterns with old patterns due to learning, as has been hypothesized previously ^19^. To test this, we explicitly modeled the encoding of a series of inputs patterns via a Hebbian plasticity rule, and set the parameters to precisely match the synaptic turnover from the fit of the network model. For simplicity, and in order to focus on the learning process itself, we made some approximations to the full network dynamics. Specifically, we modeled CA1 pyramidal cells as firing rate units with a threshold-linear activation function, and modeled inhibition by subtracting the total mean excitatory input to cells. CA3 and EC inputs were modeled as before, see Methods for details.

For the learning process, we made changes in the connections from EC and CA3 to cells in CA1 in order to store a sequence of random, sparse patterns. Once enough patterns were stored, the synaptic weight matrix reached a statistical steady state which was independent of the initial synaptic configuration. At this point we could track the drift in representation of any given pattern due entirely to the learning process itself. Specifically, we considered a rule for which potentiation of synaptic connections occurred between coactive neuronal pairs with a probability *p*_+_ while synaptic connections become depressed with a probability *p*_−_ when one of the neurons was active and the other inactive. These probabilities were allowed to be different for EC versus CA3 inputs. If neither neuron was active the synapse was not updated. Given this rule, a particular input pattern to a CA1 cell from EC or CA3 led to distinct synaptic weight structure depending on the activity of the CA1 cell itself, see Fig.5a. By virtue of this rule, connections from active presynaptic cells in EC and CA3 to active cells in CA1 became selectively potentiated for a pattern presented at time *t*, Fig.5b left. Therefore if the same input pattern was presented again, before any new learning, the resulting pattern in CA1 was highly correlated with the initial pattern, Fig.5b right. After new patterns were encoded, the synaptic weight matrix no longer strongly associated the input pattern at time *t* with the pattern of activity in CA1 at time *t*. Hence, the correlation between the activity pattern in CA1 at time *t* and that at a later time (time was measured in the number of patterns encoded) decreased.

**Fig. 5.**
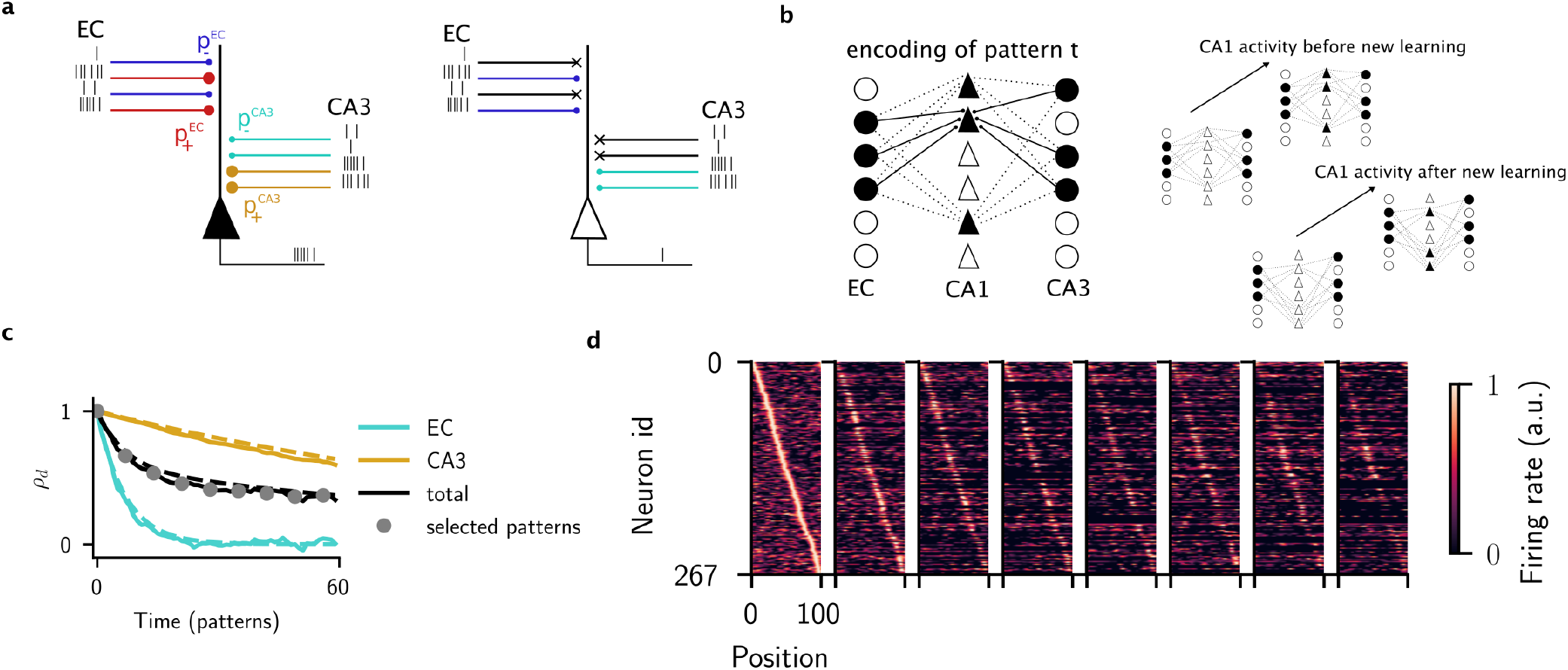
Ongoing encoding of patterns in CA1 generates representational drift. **a**, A cartoon of the plasticity rule. For a given pattern, a CA1 cell receives inputs from EC and CA3 cells and is either active (left) or inactive (right). If the pre-synaptic neuron and the CA1 cell are both active the synapse is potentiated with probability 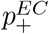 or 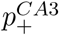 depending on the identity of the input. If the presynaptic cell is active and the CA1 cell inactive, or vice-versa, the corresponding synapse is depressed with probability 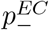 or 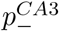. If both cells are inactive the synapse is not updated (crosses in cartoon). **b**, Cartoon of the encoding process. At time *t* a pattern of activation is chosen randomly with sparseness parameters *s*_*EC*_, *s*_*CA*1_ and *s*_*CA*3_. After the plasticity rule is applied, the connections between coactive cells will be potentiated, see left. If the input patterns are presented, the potentiated synapses will reproduce the original pattern in CA1 if there has been no new learning (right top) but not if additional patterns have been encoded (right bottom). **c**, The correlation in the number of connections to cells (degree) in CA1 *ρ*_*d*_ as a function of time as measured in patterns stored. Dashed lines are theoretical curves, solid lines are from simulation, and the circles indicate the patterns shown in d. **d**, Heatmap of the activity of CA1 cells for a particular pattern, shown at regular intervals corresponding to the circles in panel c. See methods for parameter values and model details.

In order to fit the statistics of representational drift observed in experiment, we calculated how the synaptic weight matrix changed over time as a function of the learning rates and the sparsity of the patterns. Specifically, we calculated how the correlation in the number of inputs to a cell in CA1, the degree correlation *ρ*_*d*_, decreased, depending on whether the cell participated in a given pattern or not. We chose parameters to match this decrease in correlation to that found in the network model, Fig.5c, see Methods for details. We visualized the activity of a particular stored pattern by ordering the cells according to their place field location at a particular point in time Fig.5d, leftmost panel. We then used the same ordering to observe the activity at regular intervals, corresponding to the points in time for which the drop in correlation was the same as observed in the experimental data, Fig.5d. We concluded that the observed RD in CA1 during the exploration of a familiar environment is consistent with the expected reshaping of the internal representation of space and context due to ongoing learning.

## Discussion

We have shown that the statistics of representational drift over weeks in CA1 of mice are consistent with ongoing synaptic turnover, which is random and uncorrelated with the observed patterns of activity. Specifically, a network model in which spatially untuned afferents, from EC to CA1 pyramidal cells exhibit nearly complete turnover from session to session, while correlations in spatially tuned inputs, from CA3, decay much more slowly, can reproduce an array of experimentally reported measures of RD such as the heterogeneity in single-cell stability Fig.3b, the drift in place-field centroids Fig.3h and the drop in correlation in the population vector over time, Fig.3i. Allowing for a moderate amount of spatial tuning in EC inputs does not alter this scenario, see Extended Data Fig6. The presence of two distinct time-scales of RD could be inferred directly through data analysis, Fig.1, and implemented in a simple phenomenological model to capture the statistics of active cells, Fig.2. In this simple model, input distributions were assumed to be Gaussian, with zero mean. Such input statistics arise naturally in large-scale spiking networks operating in the balanced regime ^35,37^, allowing us to map a network model onto the phenomenological model, and hence fit the data quantitatively, Fig.3.

The mechanism by which spatial tuning arises in CA1 cells in the network model tightly constrains what kind of RD can occur due to changes in inputs alone. Specifically, in the network model, CA1 cells integrate large numbers of relatively strong excitatory and inhibitory inputs, which, however, cancel in the mean, leaving an order one tuned component near threshold to spiking. Whether or not a cell is active on any given session depends on the mean level of depolarization compared to threshold. If session-to-session changes in mean input really are uncorrelated with a cell’s activity as we have assumed, then cells with elevated firing rates ought to be more stable, while low-rate cells will be more susceptible to drop out of the population code. This same mechanism also requires that the place-field width increase with increasing mean rate via an iceberg effect. We observed both of these effects in the data, Fig.4. While the experimentally observed RD was therefore broadly consistent with the proposed network mechanism, there were some important quantitative differences. Namely, the fraction of place cells in the network model was always higher than that in the data. Additionally, the within-session variability in the data was significantly larger than in the network simulations, which accounts for the discrepancy in the PV correlation for Δ*t* = 0 in the insets of Fig.3i. These differences may be due to the fact that actual CA1 cells are not solely selective to spatial location, but rather also to other behaviorally relevant features such as head direction, velocity and contextual cues ^38–41^. Modeling such selectivity requires additional inputs which we have ignored here for simplicity.

In the network model, the changes in inputs to CA1 pyramidal cells responsible for RD could arise via structural changes, changes in the patterns of presynaptic neurons (EC or CA3), or a combination of the two, see Eq.2. Chronic imaging studies have revealed a high degree of spine turnover in CA1 in vivo ^42–44^, suggesting a major role for the structural component in RD. There is also evidence that neuronal representations in CA3 are more stable than those in CA1^45,46^. We therefore chose to focus on synaptic plasticity as the main driver of RD. We modeled synaptic plasticity in the inputs to CA1 pyramidal cells explicitly via a Hebbian rule which allowed for the ongoing storage of novel patterns of activity, Fig.5. The encoding of new patterns of activity via plasticity-dependent updating of bounded synaptic weights naturally leads to the overwriting of previously stored patterns ^26,27^. The net effect of this overwriting is to change the population response in CA1 to a given input pattern, i.e. it causes drift. Importantly, the observed drift is simply the inevitable consequence of the storage of new information, and not the signature of a more complex process of adaptation of the original memory trace, nor of strategies used for learning in deep networks ^12,47^. We would argue that this mechanism for RD should be expected in any cortical circuit involved in the rapid encoding of memory. The response of cells in primary visual cortex to naturalistic movies has also been observed to undergo drift, although the response to gratings remained stable ^20,21^. This suggests that the mechanisms underlying RD in primary visual cortex may be different from what we propose here ^48^.

Other computational modeling work has also proposed that RD may arise due to plasticity-driven perturbations ^23–25^. In ^23^ the authors studied a network with prescribed synaptic weights which allow for the storage of a large number of sequential patterns. They showed that randomly perturbing the synaptic weight matrix generates RD while maintaining robust sequences. RD can also arise in a spiking network with a symmetric spike-timing dependent plasticity rule, coupled with a homeostatic mechanism ^24^. Specifically, if the initial network structure (synaptic weight matrix) exhibits clustering, ongoing plasticity allows for individual cells to leave their cluster and join a new one, all the while maintaining the clustered structure at the network level. RD occurs in a similar fashion in networks which minimize the mismatch between the similarity of pairs of input patterns and the corresponding pairs of output patterns (Hebbian/antiHebbian networks) ^25^. Namely, ongoing plasticity allows the network to explore the degeneracy in the solution space by undertaking a random walk along the manifold of equally optimal output patterns. Here we have also shown that the RD observed in CA1 of mice is not only broadly consistent with ongoing, random synaptic turnover, but even quantitatively so. An important conceptual difference between our work and previous studies is the nature of the synaptic turnover itself. While those studies showed that *already-stored* patterns of activity undergo RD in the face of ongoing plasticity, we ascribe the plasticity to the encoding of *new patterns*. Namely, we have shown that RD is consistent with the inevitable interference between patterns when learning occurs, and hence not necessarily just a consequence of noise once learning is done. The mice in the experiments we have studied explore familiar tracks, and are not exposed to explicit, task-dependent learning between sessions. Rather, the “learning” process may simply be the storage of episodes, unrelated to the exploration. The hippocampal circuit plays a central role in this type of memory ^49,50^.

*The stability paradox* We have sought here to provide a plausible network mechanism for RD in CA1, and have not addressed the fundamental paradox of how to maintain stable behavior in the face of RD. In fact, drift does not appear to adversely effect the performance of mice in a variety of memory-dependent tasks ^18,19,51^. Several previous studies have addressed how this might be possible. Firstly, it has been hypothesized that there may be a low-dimensional manifold which represents the task-relevant projection of the population response, and which is invariant to RD ^21,52,53^. In this scenario many distinct patterns of the high-dimensional population activity can have the same projection on the relevant, low-dimensional manifold. If RD does not affect this projection, i.e. it is constrained to the “null-space” of the manifold, then task-relevant variables can be stably read out. While neuronal representations undergoing RD do appear to be low-dimensional, the direction of drift has not been found to be orthogonal to this manifold in general ^9,19,54^. Alternatively, despite ongoing changes, the patterns of neuronal activity observed in-vivo retain some significant correlation from session to session ^9,10,15,18,19^. Therefore, decoders trained on a given session will perform above chance for subsequent sessions, albeit with degraded accuracy. Interestingly, the time-scale in the decay of correlation of population codes due to drift appears to be similar in hippocampus, parietal cortex and piriform cortex alike: on the order of weeks. In some studies this led to decoders reaching chance level after about a month. ^15,19^, while other groups found that a small but significant fraction of cells remained stable over the entirety of the recordings ^9,18^. In either case it remains unclear precisely how behavior remains unaffected by the observed drift. One potential solution to this is to allow for compensatory plasticity in downstream circuits so as to stabilize the readout performance ^24,52,54–56^.

Another potential solution is to seek some other higher-order population structure which remains stable in the face of ongoing RD ^21,52,53^. Work on this topic has defined stability in two very distinct ways: 1 - Resistance to drift by some sub-population of cells, i.e. a fraction of the population code does not change, or 2 - Drift occurs, but different patterns remain separable, e.g. there is a fixed correlation or geometrical relationship between them. *1 - Resistance to drift*. In the simplest case, some small but significant fraction of cells may remain in the code during the full extent of the experiment ^9,18^. It is unclear if these cells would eventually drop out of the code given enough time. Alternatively, in mouse CA1 it was found that cells which were the most “connected” to other cells in the network, as measured by the degree from the covariance matrix, formed stable subgroups ^13^. One possible confound is the fact that this degree is strongly correlated with firing rate, see Fig.S13d in ^13^, which is major predictor of stability as seen in our Fig.4. More generally, our work here suggests that the diversity in the stability of cells observed in experiment might be simply explained by heterogeneity in network properties, e.g. in the distribution of inputs. Chronic imaging experiments in mice exploring virtual tracks of up to 40 meters revealed that both the mean firing rate and the number of place fields of a given cell were broadly distributed across the population in CA1, and were highly correlated across time and environments ^15^. This property of cells, called “propensity”, may be related to differences in intrinsic excitability ^15^, although the exact mechanism responsible for the diversity in the number of place fields remains unclear. Despite the stability of propensity, the population code itself was still found to undergo RD as reported in previous studies. *2 - Drift occurs, but different patterns remain separable*. For example, in a delayed non-match to place task in which the activity of so-called “splitter” cells in the central arm of a maze predict the upcoming turn direction of the animal, the population activity always separates task dimensions despite ongoing RD ^16^. That is, although the level of activity, as well as the selectivity, of individual cells changes over time, on any given session one can decode the animal’s behavior from the population activity. This phenomenology, dubbed “heterogenous stability”, clearly does not address the problem of readout since the population activity encoding for any given task-relevant variable still changes over time. A similar phenomenon is observed in mouse hippocampus when an animal is exposed to a sequence of morphed environments which interpolate between two familiar ones ^17^. As observed previously ^7^, on any given session the neuronal representation undergoes a transition as a function of the environment along the morph axis, reminiscent of attractor-like dynamics. Nonetheless, this representation changes from session to session and hence is not stable in the sense of readout. Both of these examples are conceptually similar to earlier experiments in which the population codes for two distinct environments both exhibit RD but still remain separated over long times scales ^10^. Also, it has been discovered that under some conditions two or more distinct spatial maps may encode the exact same environment on distinct trials ^57^. Both undergo drift over time but can be reliably separated. It may seem surprising that in all of these cases the RD somehow contrives to maintain a separation in the population activity for distinct task variables or different environments. At the very least it appears to argue against RD as a blind, random process, which, intuition would have us believe, should lead to the separation between representations getting washed out. However, if distinct patterns of activity in CA1 are driven by distinct patterns of presynaptic activity, and RD is the consequence of synaptic plasticity, it is, in fact, precisely what one should observe. The reason is that the correlation in the output patterns is determined by the correlation in the input patterns and the statistics of the synaptic connectivity matrix, in particular the sparseness. Random perturbations of the connectivity matrix, e.g. due to ongoing learning, which do not change the sparseness, will generate RD but will not affect the mean correlation in the output patterns, see Extended Data Fig.8 for an illustration.

Therefore, despite the heterogeneity in the stability of cells, and despite the separation of task- or context-dependent population codes, the stability paradox remains. It may be that behavior depends on a neuronal representation which is distributed across several cortical areas, and that RD is greatly reduced in some of the them compared to those imaged up until now. In the case of the hippocampus, it is hypothesized that memories are transferred to cortical areas for long-term storage via a consolidation process which can take weeks, months or years depending on the species ^58–60^. Decades of clinical work, and experimental lesion studies in animals, indicate that many forms of long-term memory become independent of the hippocampus (although see ^61^), and hence is likely stored in neocortical circuits ^59,62,63^. Computational models of distributed memory systems in which fast learning (and hence fast forgetting) circuits drive plasticity in slower-learning downstream circuits in a multi-layered framework, show qualitatively enhanced memory capacity compared to single-area models ^28^. Such models leverage a hierarchy of time-scales of synaptic plasticity to allow for both fast encoding as well as long lifetimes ^28,64,65^. A hallmark of a model in which the timescale of plasticity is spatially distributed across cortical areas, fastest for hippocampus, and increasingly slower for downstream cortical areas, is a concomitant array of timescales for RD. Only future experiments in which the activity of neuronal populations across several cortical areas are recorded simultaneously over long periods of time will reveal if RD acts on distinct timescales in different brain areas.

## Methods

### Data analysis

We analyzed previously published data ^10^. In the experiment, mice repeatedly explored two familiar environments (one in the morning, the other in the afternoon) over the course of two weeks, with imaging sessions every other day (8 total sessions). We separately analyzed leftwards vs rightwards running epochs for each environment, and then pooled the data together. In total, we analyzed data from 5 mice. Of the five mice, two have only one spatial map for each environment, while three have multiple spatial maps for each environment, as previously analyzed ^14^. Unless otherwise specified, all the analysis was performed separately for each map, and then results were pooled together.

Calcium events and place field maps were extracted as in the original paper ^10^. Briefly, the linear tracks were divided into 24 bins, each 4cm in length. For each spatial bin, the total number of events in each session was extracted, together with the total time-occupancy of each bin. The event rate map is then calculated by dividing the total number of events per bin by the bin occupancy. The two bins at each extreme of the track were excluded from the analysis to limit reward delivery effects.

In Fig. 1a-b, the rate maps follow two different types of normalization: in Fig. 1a, the event rate map for each cell was normalized by the maximal firing rate over all sessions. In Fig. 1b, the event rate in each session/bin as normalized according to the maximal value within each session.

### Rate and tuning correlation

To compute rate correlations across sessions, we defined a rate population vector **r**_**t**_ ∈ ℛ^*N*^. *N* was the total number of neurons, and each entry of the vector was the mean firing rate of each cell in session *t*. We then computed the Pearson correlation coefficient to quantify the rate similarity between two sessions, as in Fig. 1d-f.

To quantify the similarity of spatial tuning across session, we defined a Tuning population vector of length *N* × *n*, where *n* is the number of bins of the linear track. The vector then contained the rate of each cell in each spatial bin, normalized in such a way that the sum of the rates for each cell over the different spatial bins was constant in all sessions. We then computed the Pearson correlation coefficient of such vectors for pairs of session to quantify the tuning similarity. The procedure ensured that changes in mean firing rates of cells from one session to the next would not affect the tuning correlation.

### Place fields

To be considered a place cell in a given session, we employed the following procedure. First, for each cell, we computed the spatial information per spike as :

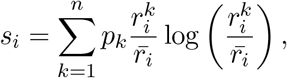

where *i* was the cell index, and the sum runs over all *n* spatial bins, 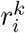 was the event rate of cell *i* in bin *k*, 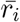 was the mean event rate over all bins, and *p*_*k*_ was the occupancy of bin *k* (the fraction of time spent in bin *k*). *s*_*i*_ was then the spatial information (measured in bits/event) of cell *i*. Then, we generated surrogate data by shuffling the position of the animal with respect to the time of the calcium events, and calculated the spatial information for each cell and each shuffle. We then compared the value of the spatial information of each cell to the null distribution generated with the surrogate data. If the value was larger than the 95th percentile of the null distribution, than the cell was defined as a place cell in that session.

The place field width of each cell was defined as the number of contiguous bins where the event rate is larger than 50% of its maximal value over all the bins.

### The statistical model

We modeled cells in CA1 as binary units which received inputs from two sources, CA3 and layer III of entorhinal cortex EC. The total input to a cell *i* at time *t* (time measured in session) from CA3 was written 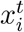 and from EC 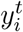. Both inputs were Gaussian random variables with zero mean and variances 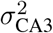 and 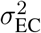 respectively. A neuron *i* was active at time *t* if its total input 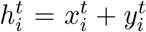 exceeded a threshold *θ*, and was otherwise silent. Specifically, the activity was written as 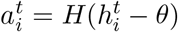, where *H*(*x*) = 1 if *x* > 0 and *H*(*x*) = 0 if *x* ≤ 0 is the Heaviside function.

To model RD we allowed for the inputs to change over time according to:

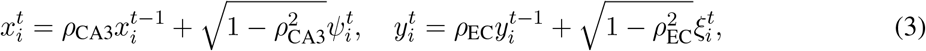

where the autocorrelations *ρ*_CA3_ and *ρ*_EC_ ∈ [0, 1] and 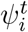 and 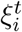 were Gaussian random variables with mean zero and standard deviation *σ*_CA3_ and *σ*_EC_ respectively. The update rules Eqs. (3) ensured that the distribution of inputs were stationary. Inhibition is implicitly assumed to have the effect of subtracting the mean of both inputs so that they are centered at zero.

The statistical model had four parameters once we rescaled them by the standard deviation of the inputs from EC: the ratio *σ*_CA3_*/σ*_EC_, the rescaled threshold *θ/σ*_EC_ and the autocorrelations *ρ*_CA3_ and *ρ*_EC_. Note that, the input dynamics Eqs. (3) could be formulated in continuous time as an Ohrnstein-Uhlenbeck processes, as shown in the Supplementary Material. In the continuous formulation allowed us to calculate the time constant of the decay in correlation of the inputs analytically, yielding 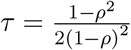.

According to the definitions above, the state of the network was defined by a vector 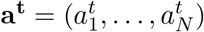 where *N* was the total number of neurons. *Fraction of active cells:* The fraction of active cells can be calculated analytically for the statistical model. The probability that a cell with a given input *y* is active can be written Pr(*y* > *θ* − *x*). Integrating this probability over all possible values of *y* gives the likelihood of any cell to be active, or the fraction of active cells, *f*_*a*_. This fraction is therefore

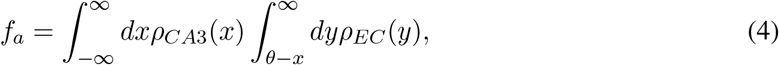

where 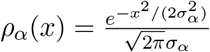

### Fit of the statistical model

To fit the statistical model, we only considered mice with a single map per environment (two mice in total), since with multiple maps the distribution of the number of sessions in which each cell was active and the survival fraction are not well defined (with multiple maps not all maps are visited in all sessions). We considered data from these two mice (N=1649 total recorded cells). The statistics used to fit the model were the distribution of sessions each neuron is active, the survival fraction (probability an initially active cell continued to be active on subsequent sessions), the population activity overlap between different sessions, and the fraction of active cells in each session. To fit the data, we adjust the four free parameters of the model using least-square optimization. Specifically, to produce Extended Data Fig. 2, we discretize the (*ρ*_*CA*3_, *ρ*_*EC*_) space on a 20 × 20 grid, and for each position on the grid, run the Scipy ^66^ implementation of a Basin-hopping optimization algorithm minimizing the sum of the squared residuals between numerical simulations of the statistical model, and the experimental data. For the numerical simulations of the model, we consider *N* = 20000 neurons.

### Network model description

CA1 was modeled as a network of integrate-and-fire excitatory (E) and inhibitory (I) neurons with I-I, E-I and I-E connections, but no recurrent excitation, as prescribed by anatomical constraints. CA3 cells are modeled as Poisson neurons a fraction of which were spatially modulated. They projected onto both excitatory and inhibitory CA1 neurons. Additionally, CA1 neurons received excitatory inputs from a layer of non-spatial Poisson neurons (from layer III of EC). A schema of the network model is illustrated in Fig. 3a. Detailed equations and parameters are given in appendix.

### CA3 place fields and non-spatial inputs

CA3 neurons were modeled as Poisson neurons. In any given environment, a fraction *f*_*CA*3_ of CA3 neurons were active. Of the active neurons, a fraction *f*_*s*_ of the population had a spatially modulated firing rate, while the remaining fraction had a constant firing rate. For simplicity, we considered a ring topology, so that the spatial position of a virtual animal was parametrized by an angle *ϕ* ∈ [− *π, π*]. The firing rate of spatially selective neurons was modulated according to a Von Mises distribution:

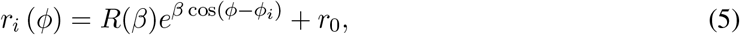

where *ϕ*_*i*_ was the center of the place field of neuron *i, r*_0_ a baseline firing rate, and 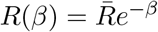 where 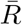 was a constant that set the maximal firing rate. The parameter *β* determined the sharpness of the place fields.

The layer of neurons providing non-spatial inputs were modeled as Poisson neurons with constant firing rate *ν*. Also for this layer, only a fraction *f*_*EC*_ of the population was active in any given environment.

### Connectivity matrices

Connectivity matrices for all subtypes of connections within the CA1 populations were random and sparse. Each neuron had a probability of connection to other neurons in the respective subpopulations equal to *α*_*i,l*_ = *K*_*i,l*_*/N*_*l*_, where *N*_*l*_ was the number of neurons of the *l*th population, *l* ∈ {*E, I* }, and *i* ∈ {*E, I*}. If not specified otherwise, we fixed the in-degree of CA1 pyramidal cells from interneurons to be fixed and equal to *K*_*I*_ (all pyramidal cell receive the same amount of inhibitory inputs). On average, each CA1 cell received projections from *K*_*i*,{*E,I*}_ neurons. The connectivity between the layer of non-spatial Poisson neurons and CA1 is also random and sparse with connection probability *α*_*EC*_. In the main text, we consider a uniform connection probability also for the CA3 → CA1 projections; in appendix we discuss how to consider a phase bias in such connections.

Throughout, we assume all connection probabilities are the same and equal to *α* = 0.125.

### Input correlations in the network model

To implement the session-to-session changes in spatial and non-spatial inputs provided by Eqs. (3) in the network model, we needed to calculate the autocorrelation of inputs where changes may occur either due to changes in the input firing pattern, or in the connectivity matrices themselves. The general expression for such inputs is

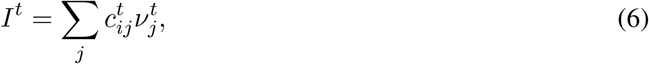

where 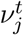 is the firing rate of the input neuron *j* at time *t*, 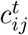 is the connectivity matrix element at time *t*, and we omitted conductances and time constants. In appendix we provide detailed calculations. The expression one finds for the autocorrelation is

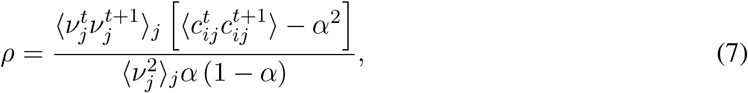

where *α* is the connection probability to the input layer. In general then, the level of correlation depends on the correlation in the input firing patterns, and on the degree of synaptic plasticity. Note that in order to obtain completely uncorrelated inputs from one session to the next, one must have 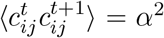, which implies 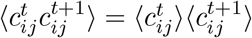, i.e. complete rewiring from one session to the other. Assuming completely uncorrelated input firing patterns from one session to the next results in:

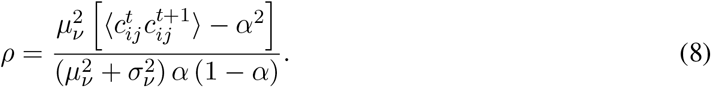

On the other hand, if we assume that the input firing patterns are the same over sessions, we have:

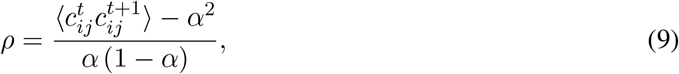

which, as shown in Appendix, can be written as:

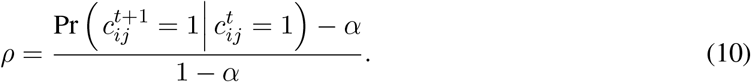

In the following, for both spatial and non-spatial inputs, we assumed that the input firing rates were constant from one session to the next. In this case, the fraction of connections rewired from one session to the next depending on the autocorrelation *ρ* is shown in Fig. Extended Data Fig. 7.

### Variance of spatial and non spatial inputs

The average current from either the spatial or non-spatial inputs layer had the form:

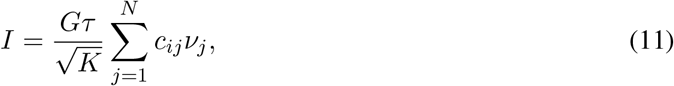

where we have scaled the synaptic weights as 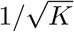. The expected value of the input is therefore

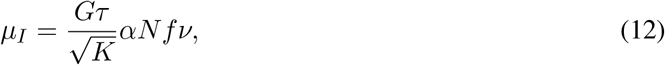

where *α* is the connection probability, *f* is the fraction of active pre-synaptic cells, and *ν* is their mean rate. If we define the normalization factor as the mean number of active inputs *K* = *αf N*, we have 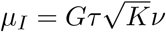. The variance of such current, using the results from appendix, was given by

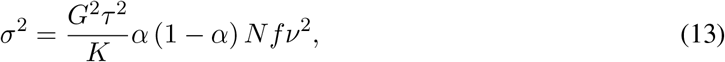

where we neglected the intrinsic variability of the Poisson process (which goes to zero as Δ*t*^−1^). Again using the definition of the normalization factor *K*, we have

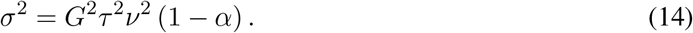

Given the variances ratio obtained fitting the statistical model 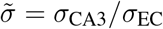, we can then fix the firing rates/synaptic weights of CA3 and EC inputs via the relation:

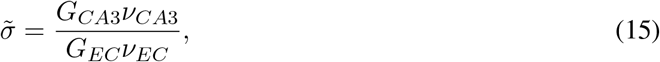

since both the time constant and the connection probabilities are the same for the two layers. We then fix *ν*_*CA*3_ = *ν*_*EC*_ (same average rate of the two layers), and set 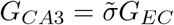.

### Fit of network simulations to data

In network simulations the virtual animal runs at a constant speed of *v* = 12 cm/s over a circular track of length *L* = 84 cm. The position of the animal on the track is parametrized with a phase *ϕ* ∈ [− *π, π*]. Analogously to the experiment, we simulate 8 sessions each consisting of 20 laps on the track (i.e. one session lasts 140 seconds of simulation time). For place fields analysis, the track was divided into 20 bins each of 4.2 cm length.

Given the analytical formulas derived in the previous section we were able to match both the variances of the input distributions as well as their autocorrelations to the values from the fit of the statistical model. The means of the input distributions were not zero, as in the statistical model, but the network operated in a balanced regime in which currents from inhibitory interneurons cancelled the mean excitatory drive to cells in the mean, leaving their membrane potential near threshold to spiking. Therefore, fitting the variances and correlations alone set the network at a working point in which the statistics of RD were close to the statistical model, and hence the data. We then made slight changes to parameters by hand in order to improve the fit. In order to fit the population vector correlation we needed to compare to the calcium event rate in the data. We did this by applying an exponential kernel with time constant *τ*_*c*_ = 500ms to the spike train from each cell in the spiking network. A calcium event was detected whenever the smoothed signal crossed a threshold *θ*_*c*_, and imposed a minimum inter-event interval of 500ms. The threshold value which minimized the mean squared error of the fit was *θ*_*c*_ = 0.16837.

### Plasticity model

We modeled a population of CA1 cells as linear threshold units, i.e. the activity of cell *i* was given by *r*_*i*_ = [*I*_*i*_ − *θ*]_+_, where *I*_*i*_ was the total input, *θ* was the threshold, and [*x*]_+_ = *x* if *x* > 0 and is zero otherwise. Time was taken to be discrete. At each time point random patterns of activity were imposed in EC, CA1, and CA3, with sparseness *f*_*EC*_, *f*_*CA*1_ and *f*_*CA*3_ respectively. Given these patterns, plasticity occurred in the synaptic weights *EC* → *CA*1 and *CA*3 → *CA*1 according to the following Hebbian rule. If pre- and postsynaptic cells were coactive, then the synapse was potentiated with probability 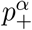, while if one cell was active and the other inactive, then the synapse was depressed with probability 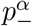, where *α* ∈ {*EC, CA*3}. If neither cell was active no change was made. Synapses were taken to be binary, i.e. a synapse from cell *j* in population *α* to cell *i* in CA1 was written 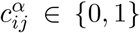. After a sufficient number of time steps the synaptic weight matrix reached a stationary state in which the statistics did not change further (although plasticity was ongoing) and were independent of the initial state of the matrix. Active cells in EC had a constant firing rate *r*_*EC*_ while active cells in CA3 were spatially modulated according to a von Mises distribution as in Eq.5. The parameters used to generate Fig.5c,d were: *f*_*EC*_ = *f*_*CA*1_ = *f*_*CA*3_ = 0.5, 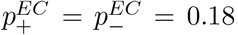, 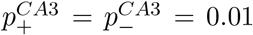 *N*_*EC*_ = *N*_*CA*1_ = *N*_*CA*3_ = 1000 and *θ* = 0.3. For the von Mises rates the parameters were *r*_0_ = 5, 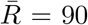, *β* = 19. *r*_*EC*_ is chosen such that the ratio *σ*_*CA*3_*/σ*_*EC*_ = 1.16 in order to match the fit from the statistical and network models. The initial condition for the connectivity matrix was random with connections probability *p* = 0.33. We tracked the 100th pattern stored, and calculated the decrease in the correlation of the degree distribution of cells (Fig.5c), and plotted snapshots of the activity in CA1 starting at time *t* = 140 and every 7 time steps for eight “sessions”, Fig.5d. Neurons were considered inactive if their mean firing rate was less than 0.1. The analytical calculation of the drop in correlation of the degree over time (dashed lines in Fig.5c) can be found in the Appendix.

## Acknowledgements

AR and FD thank Yaniv Ziv and Alon Rubin for the experimental data as well as for many helpful and enlightening discussions. AR and FD also thank Klaus Wimmer, Alex Hyafil and Jose M. Esnaola for helpful discussions and feedback. AR acknowledges “Retos” project RTI2018-097570-B-100 from the Ministry of Science and Innovation of the Spanish Government, Flag-Era project from the EU for the Human Brain Project HIPPOPLAST (Era-ICT code PCI2018-093095), “Red de Investigación” RED2018-102323-T from the Ministry of Science and Innovation of the Spanish Government. This work is supported by the Spanish State Research Agency, through the Severo Ochoa and Maria de Maeztu program for Centers and Units of Excellence in R&D (CEX2020-001084-M). We thank CERCA Program/Generalitat de Catalunya for institutional support.

