## Supplementary Figs and Text for "Network mechanisms underlying representational drift in area CA1 of hippocampus"

### Supplementary Figures

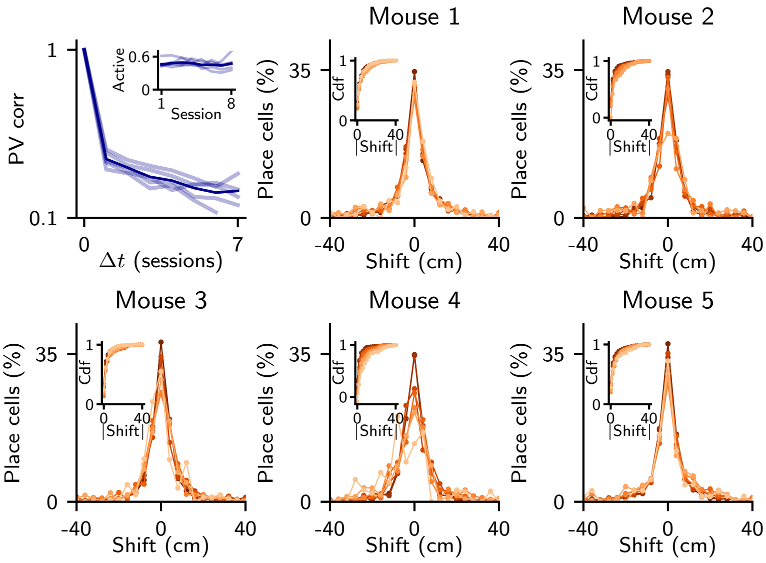


**Extended Data Fig. 1 | Population vector correlation and centroid shifts for individual mice. The dark purple line indicates the mean of the PV correlation, while the lighter lines are the PV correlation for the individual animals. The shading for the centroid shifts go from darker to lighter for an increasing number of sessions elapsed (from 1 to 7 sessions).**

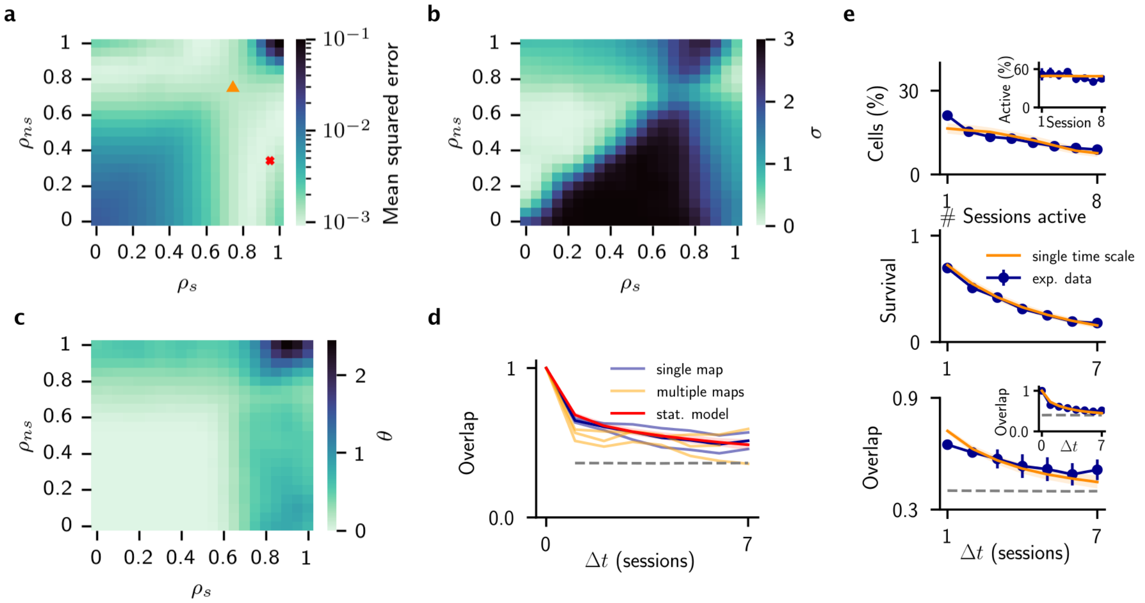


**Extended Data Fig. 2 | Fit of the data. a,** Colormap showing the mean squared error of the fit in the $(\rho_{\mathrm{CA3}}, \rho_{\mathrm{EC}})$ plane. Red cross indicates parameters chosen for Fig.2, orange triangle parameters of panel **e**. **b,c** Colormaps showing parameters $\sigma$ and $\theta$ for the solutions found in panel **a**. **d** Overlap over time for mice with a single map per environment (blue, also shown in Fig.2) and multiple maps per environment (yellow). **e**, Simulations of the statistical model where we impose both inputs to evolve on the same time scale (dark triangle in panel **a**).

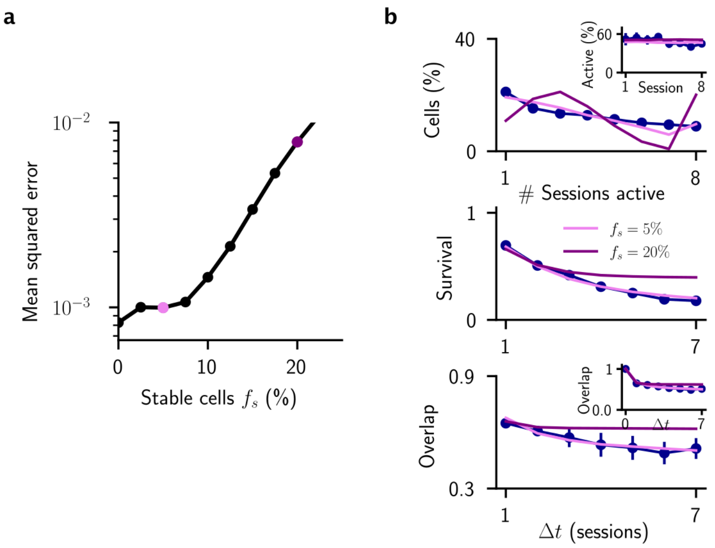


**Extended Data Fig. 3 | Fit of the model with stable fraction of cells. a,** Error of the fit as a function of the percentage of stable cells in the model. Pink and purple dots correspond to simulations of panel **b**. **b**, Best fit of the model with 5 % (pink) and 20 % (purple) of stable cells.

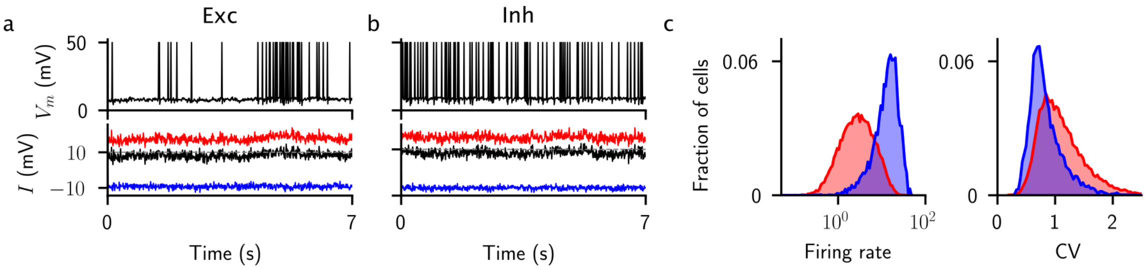


**Extended Data Fig. 4 | The network operates in the balanced regime. a,** Membrane voltage (top) and excitatory (red), inhibitory (blue) and total (black) currents (bottom) into an example excitatory neuron over one lap on the track (lap time: 7s). Excitatory and inhibitory currents balance, so that the total input is right below threshold. Shading indicated the place field of the cell. **b**, Same as **a**, but for an example inhibitory cell. **c,** Distribution of mean firing rates (left) and ISI coefficient of variations (right) for excitatory (red) and inhibitory (blue) cells. Data pooled across all sessions. Only cells with >10 spikes per session are considered.

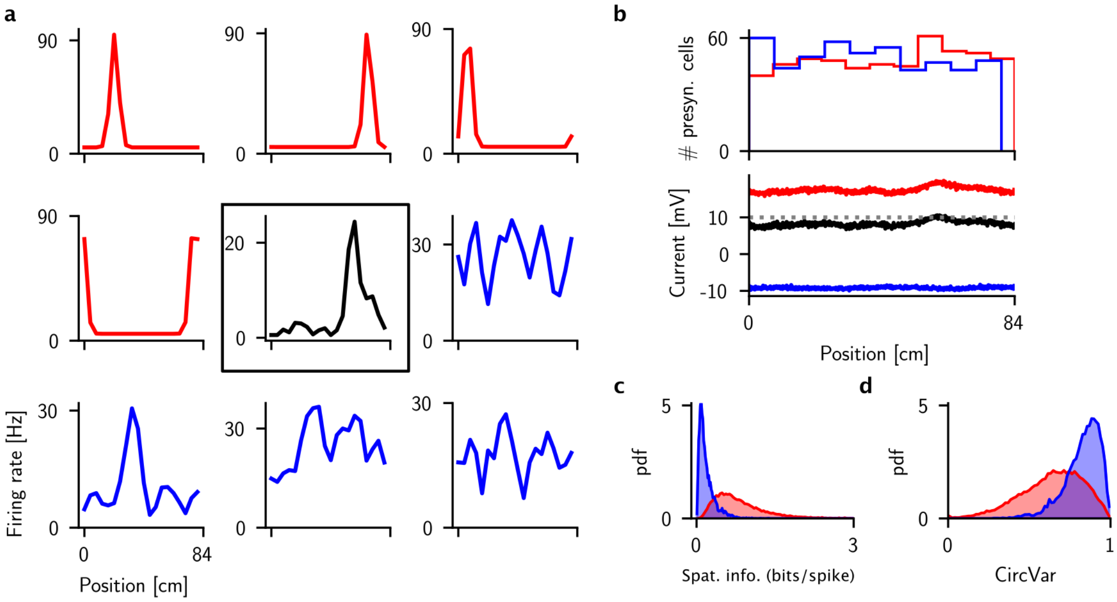


**Extended Data Fig. 5 | Emergence of selectivity in the network. a,** Tuning curve of the same cell as in panel a of Fig.4 (black), and of 8 of its excitatory (red) and inhibitory (blue) neurons. Only tuned excitatory neurons are considered. **b,** Distribution of place field positions of all inputs to the cell highlighted in panel **a** (top), and the corresponding mean excitatory (red), inhibitory (blue) and total (black) synaptic input.

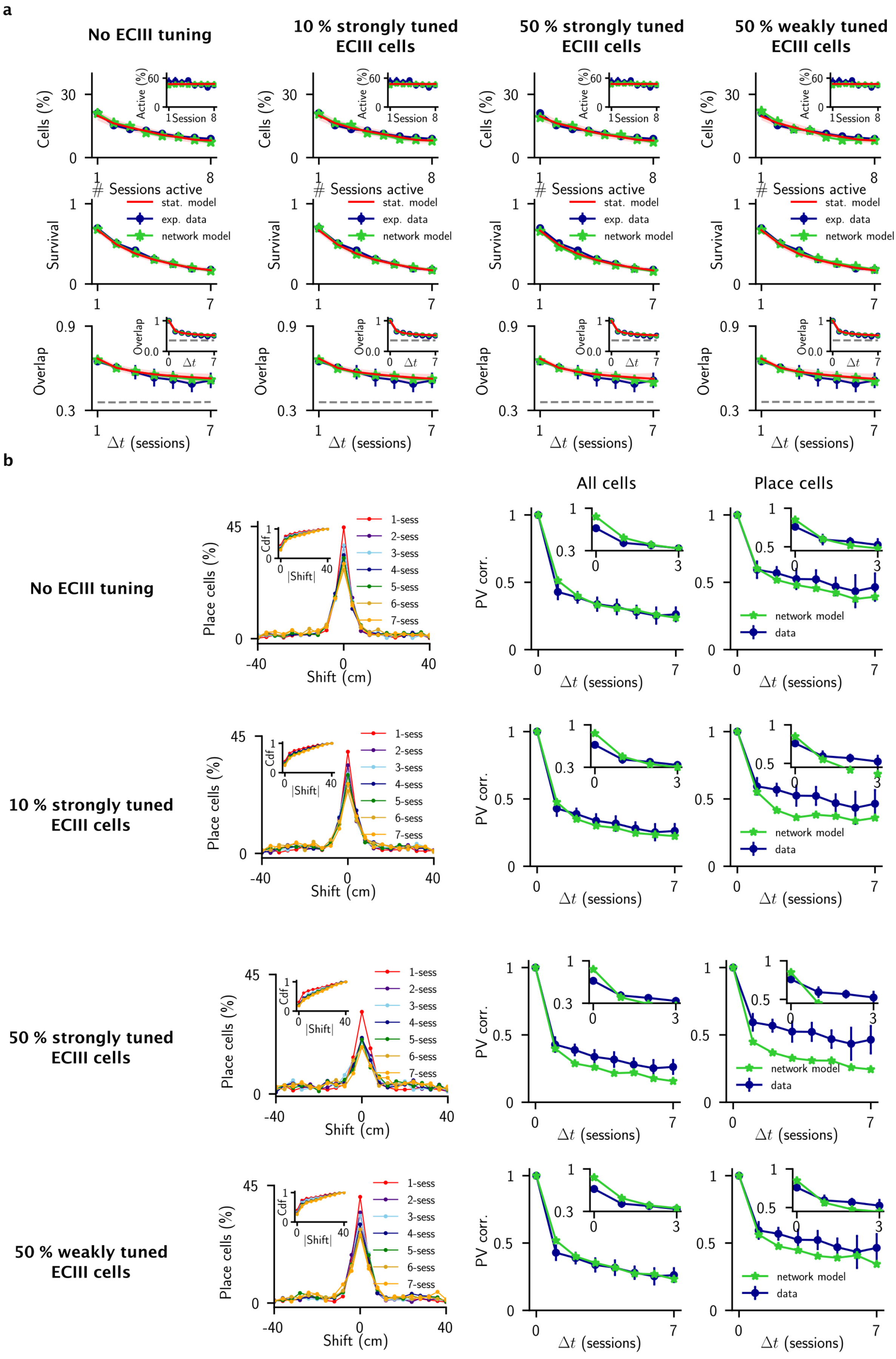
a
No ECIII tuning
10 % strongly tuned ECIII cells
50 % strongly tuned ECIII cells
50 % weakly tuned ECIII cells
Cells (%)
Active (%)
Session
### Sessions active
Survival
stat. model
exp. data
network model
Overlap
Δt (sessions)
b
No ECIII tuning
10 % strongly tuned ECIII cells
50 % strongly tuned ECIII cells
50 % weakly tuned ECIII cells
Place cells (%)
Cdf
|Shift|
Shift (cm)
1-sess
2-sess
3-sess
4-sess
5-sess
6-sess
7-sess
All cells
Place cells
PV corr.
network model
data
Δt (sessions)

**Extended Data Fig. 6 | The effect of tuning in EC on RD. a,** Adding tuning to EC does not alter the statistics of active cells. **b,**: Increasing the tuning in EC cells leads to increased diffusion in the position of place cell centroids (decreased peak in histogram of centroids shifts), and a sharper drop in PV correlation of place cells. This affect is most prominent when cells in EC are tuned according to a von Mises distribution, as in CA3, i.e. "strongly tuned". Weaker tuning, of a cosine type, does not lead to significant effects, even if the fraction of tuned cells is large, see bottom row. All parameters are as in Fig.3.

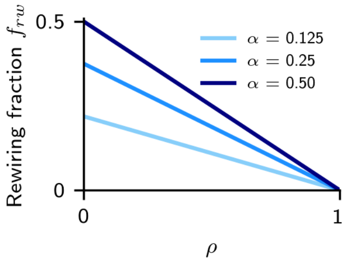


**Extended Data Fig. 7 | Fraction of connections changing from one session to the next $f_{rw}$ as a function of the correlation** $\rho$. The value is given by the formula $f_{rw} = 2\alpha\,(1-\alpha)\,(1-\rho)$.

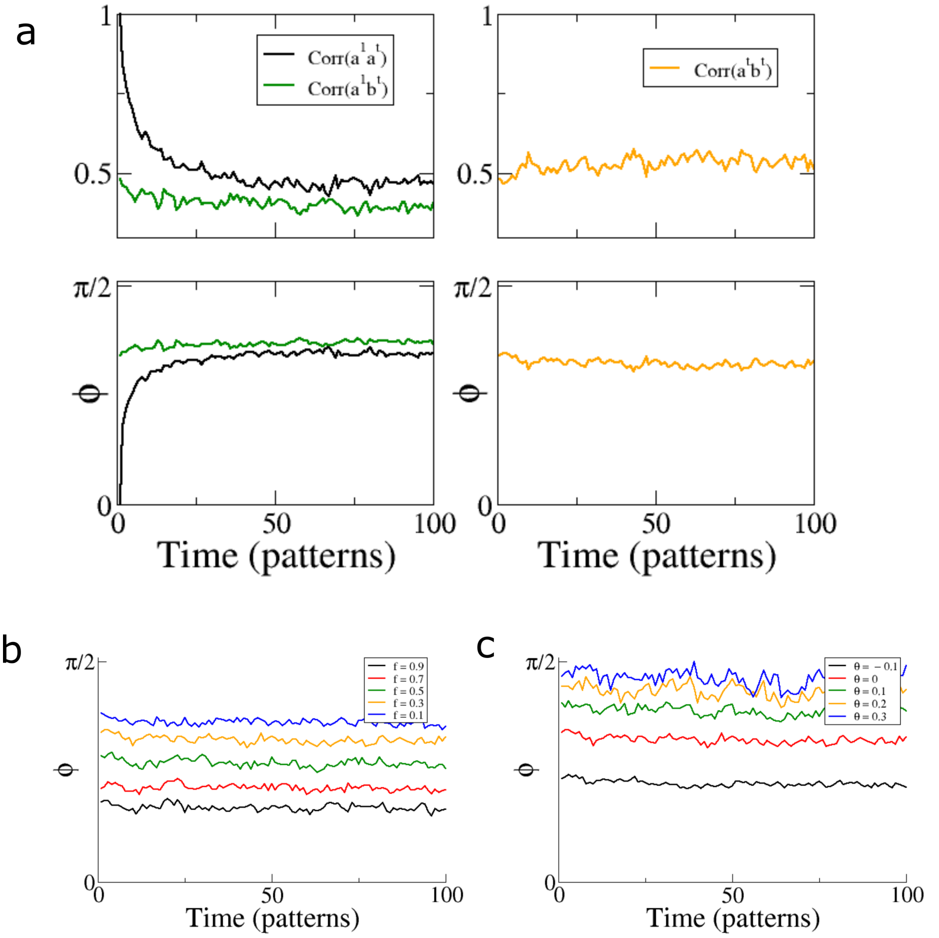


**Extended Data Fig. 8 | Population codes for distinct "contexts" remain separable despite RD caused by random, ongoing plasticity. a** The change in correlation and angle between two patterns of activity over time. The model consists of a layer of binary neurons which receive input from an upstream layer of neurons, also binary, and each of size $N$. Feedforward connections are random and sparse, with connection probability $\alpha$. Two distinct inputs patterns of sparseness $f$ are presented, which generate two binary output vectors, $a$ and $b$ once the input to a postsynaptic neuron is compared to a threshold $\theta$. At each point in time synapses are subjected to a depression (pruned) with probability $p_-$, or a potentiation (set to one) with probability $p_+ = p_-\alpha/(1-\alpha)$. Left: top - The correlation between the initial state of vector $a$, called $a^1$ and vector $a$ at later times (black), and the correlation between $a^1$ and vector $b$ at later times (green). bottom - The angle between the two vectors $\phi$. Right: The correlation and angle between the vectors $a$ and $b$, but always measured at the same point in time. Note that the correlation (or the angle) does not change. **b** The angle between the vectors $a$ and $b$ over time for different levels of sparseness of the presynaptic pattern of activation $f$. In the limit of $f = 1$ the presynaptic patterns become identical and so the angle between $a$ and $b$ goes to zero, while in the limit $f \to 0$ the patterns become orthogonal. **c** The angle between the vector as a function of the threshold of the postsynaptic neurons. The larger the threshold, the sparser the activity: for $\theta = 0$ the sparseness is one half, while for $\theta = 0.3$ it is about 0.05. Parameters, unless noted otherwise, are: $N = 1000$, $\alpha = 0.1$, $p_- = 0.1$.

### Supplementary Information

#### Input dynamics as an Ornstein-Uhlenbeck process

The update rules Eq.3 can be rewritten, in continous time, in this form:

$$\epsilon\frac{dz}{dt} = \sqrt{1-\rho^2}\sigma\eta(t) - (1-\rho)\,z(t), \tag{16}$$

where $\langle\eta(t)\rangle = 0$ and $\langle\eta(t)\eta(s)\rangle = \delta(t-s)$.
Eq. (16) is a Ornstein-Uhlenbeck with drift coefficient $\mu = (1-\rho)/\epsilon$ and diffusion coefficient $D = \sigma^2\left(1-\rho^2\right)/(2\epsilon^2)$. The associated Fokker-Planck equation for the probability distribution of the input $P(z,t)$ is then:

$$\frac{\partial P(z,t)}{\partial t} = \mu\frac{\partial\,(zP)}{\partial z} + D\frac{\partial^2 P}{\partial z^2}. \tag{17}$$

The stationary solution of equation (17) is a gaussian distribution centered at zero with variance

$$\sigma^2 = \frac{D}{\mu} = \frac{\sigma^2\left(1-\rho^2\right)}{2\epsilon\,(1-\rho)}. \tag{18}$$

We therefore set $\epsilon = \left(1-\rho^2\right)\left[2\,(1-\rho)\right]^{-1}$ so that, if the initial distribution of $z$ is $\mathcal{N}(0,\sigma)$ the process is stationary. Eq. (16) is then

$$\frac{dz}{dt} = \sqrt{2D}\eta(t) - \mu z(t), \tag{19}$$

where

$$\mu = \frac{2}{(1-\rho^2)}\left[1-\rho\right]^2 \quad D = \sigma^2\mu. \tag{20}$$

In the limit $p \to 0$, we have $dz/dt = 0$. When $p$ goes to 1, $dz/dt = 2(\sigma\eta - z)$.
The covariance of a process of the type Eq.(16) is given by:

$$\langle z(t)z(0)\rangle = \sigma^2 e^{-t/\tau}, \tag{21}$$

where we defined $\tau = \mu^{-1}$. Note that $\lim_{\rho\to 0}\tau = 1/2$ and $\lim_{\rho\to 1}\tau = \infty$: when there is no plasticity, the autocorrelation decay time is infinite.

#### Network equations

Each model neuron follows the LIF dynamics:

$$\tau\dot{V}_i^l = -V_i^l + \tau I_i^l \tag{22}$$

with $i = 1,\ldots,N_l$ and $l \in \{E, I\}$. $I_i^l$ is the synaptic current. The neuron emits a spike when it crosses $V_{\text{th}} = 10$ mV, where it is then resetted to $V_r = 0$ mV.
The synaptic current to excitatory neurons is:

$$I_i^E = \frac{g_s}{\tau_s}\sum_{j=1}^{N_{\text{CA3}}} c_{ij}^{E,s}\sum_k e^{-\left(t-t_{j,k}^s\right)/\tau_s} - \frac{g_{EI}}{\tau_s}\sum_{j=1}^{N_I} c_{ij}^{E,I}\sum_k e^{-\left(t-t_{j,k}^I\right)/\tau_s} + \frac{g_{ns}}{\tau_s}\sum_{j=1}^{N_{\text{EC}}} c_{ij}^{E,ns}\sum_k e^{-\left(t-t_{j,k}^{ns}\right)/\tau_s} \tag{23}$$

where $g_{s,EI}$ are the feedforward and E ← I synaptic conductances, $t^l_{j,k}$ are the times of the $k$th action potential of neuron $j$ of the $l$ population, and $C^{l,r}$ the connectivity matrices.
The synaptic current to inhibitory neurons is:

$$I^I_i = \frac{g_{IE}}{\tau_s}\sum_{j=1}^{N_E} c^{I,E}_{ij}\sum_k e^{-\left(t-t^E_{j,k}\right)/\tau_s} - \frac{g_{II}}{\tau_s}\sum_{j=1}^{N_I} c^{I,I}_{ij}\sum_k e^{-\left(t-t^I_{j,k}\right)/\tau_s} +$$
$$\frac{g_{Is}}{\tau_s}\sum_{j=1}^{N_{\mathrm{CA3}}} c^{E,s}_{ij}\sum_k e^{-\left(t-t^s_{j,k}\right)/\tau_s} + \frac{g_{Ins}}{\tau_s}\sum_{j=1}^{N_{\mathrm{EC}}} c^{E,ns}_{ij}\sum_k e^{-\left(t-t^{ns}_{j,k}\right)/\tau_s} \quad (24)$$

Unless otherwise stated, we set $g_{Is} = g_s$ and $g_{Ins} = g_{ns}$. To set the network in the so called balance state, the synaptic weights are scaled according to:

$$g_l = \frac{G_l}{\sqrt{K}}, \quad (25)$$

where $K$ is the average number of active inputs to each subpopulation, and $G_l$ is indepedent of $K$

#### Network parameters

The membrane time constant is set to $\tau = 10$ ms, and the synaptic time constant $\tau_s = 5$ ms. The synaptic weights of CA1 recurrent connections are $G_{EI} = 2.8$ mV, $G_{IE} = 2.0$ mV, $G_{II} = 2.0$ mV[1]. The other synaptic weights (that determine the variances of spatial and non-spatial inputs) are set in the following considering the fit of the statistical model. Parameters of the CA3 place fields are $\beta = 19.85$, $r_b = 5$ Hz, $\bar{R} = 90$ Hz.
Unless specified otherwise, we consider $N_E = 4000$ and $N_I = 1000$ CA1 cells. The connections probability is the same for all connectivity matrices and equal to $\alpha = 0.125$. The sparsity of the CA3 and EC layer are set respectively to $f_{CA3} = 0.5$ and $f_{EC} = 0.5$. We wish to have about the same number of excitatory CA1, CA3 and EC active cells, so that we set $N_{CA3} = N_E/f_{CA3} = 8000$ and $N_{EC} = N_E/f_{EC} = 8000$. The fraction of tuned CA3 inputs $f_s$ is set to $f_s = 0.5$, unless specified otherwise.

#### Input correlations in the network model

In this appendix, we calculate the amount of correlations of the inputs expected in the network model, for the same or different environments. In the network model, average synaptic inputs to a cell $i$ in environment $A$ can be written in the form:

$$I^A_i = \sum_j c^A_{ij}\nu^A_j, \quad (26)$$

where we omitted conductances and time constants. $c^A_{ij}$ is the matrix element during visits of environment $A$ (if visits do not occur at the same time plasticity may have occurred), and $\nu^A_j$ is the firing rate of input neuron $j$ in environment $A$.
In general, the Pearson correlation of inputs from different environments can be written as:

$$\rho = \frac{\langle I^A I^B\rangle - \mu_A\mu_B}{\sigma_A\sigma_B}, \quad (27)$$

where we have defined $\mu_{A,B} = \langle I^{A,B}\rangle$ and dropped the index $i$ for simplicity. Note that the bracket averages are all over the index $i$ (i.e. we consider one specific realization of the input neurons rate $\nu$).

[1]Note that these weights are then rescaled according to Eq.25.

The expected values take the form:

$$\mu_{A,B} = \sum_j \langle c_j^{A,B} \nu_j^{A,B} \rangle = \alpha \sum_j \nu_j^{A,B}, \tag{28}$$

where for the second equality we have assumend that the connection probability $\alpha$ in the two environments is the same. On the other hand, we have for the expected value of the product of the inputs:

$$\langle I^A I^B \rangle = \sum_j \sum_l \langle c_j^A c_l^B \nu_j^A \nu_l^B \rangle = \sum_j \langle c_j^A c_j^B \rangle \nu_j^A \nu_j^B + \sum_{j \neq l} \langle c_j^A c_l^B \rangle \nu_j^A \nu_l^B. \tag{29}$$

We then have

$$\langle I^A I^B \rangle = \sum_j \nu_j^A \nu_j^B \langle c_j^A c_j^B \rangle + \alpha^2 \sum_{j \neq l} \nu_j^A \nu_j^B. \tag{30}$$

For the within-environment variances, we have:

$$\left(\sigma^{A,B}\right)^2 = \sum_j \sum_l \nu_j \nu_l \langle c_j c_l \rangle - \mu^2 = \alpha \sum_j \nu_j^2 + \alpha^2 \sum_{j \neq l} \nu_j \nu_l - \alpha^2 \sum_{j,l} \nu_j \nu_l, \tag{31}$$

where we dropped the indeces A,B for simplicity and substituted the previously found expression for $\mu$. Adding and removing the element where $j = l$ in the second term gives:

$$\sigma^2 = \alpha \left(1 - \alpha\right) \sum_j \nu_j^2. \tag{32}$$

Using the same trick in the numerator of adding and removing one element from the sum, we get for the correlation:

$$\rho = \frac{\sum_j \nu_j^A \nu_j^B \left( \langle c_j^A c_j^B \rangle - \alpha^2 \right)}{\alpha \left(1 - \alpha\right) \left[ \sum_j \left( \nu_j^A \right)^2 \right]^{1/2} \left[ \sum_j \left( \nu_j^B \right)^2 \right]^{1/2}}. \tag{33}$$

If we now consider that the input layer is very large, then all sums over $j$ can be rewritten in terms of mean values. If we further assume that the input patterns are drawn from the same distribution for the different environments, then we have:

$$\rho = \frac{\langle \nu_j^A \nu_j^B \rangle_j \left[ \langle c_j^A c_j^B \rangle - \alpha^2 \right]}{\langle \nu_j^2 \rangle_j \alpha \left(1 - \alpha\right)}, \tag{34}$$

where $\langle \cdot \rangle_j$ stands for the average over the input layer.
Eq.(34) is the general form of the inputs considering different environments/directions or the same environment at different times. We now consider three separate cases:

1. Visits of the same environment at different times (same inputs firing rates but different connectivity matrices)

2. Partially correlated environments at approximately the same time (different input firing rates but constant connectivity)

3. Orthogonal environment at different times.

**Case 1.** If the environment is the same, at least for the spatial input we can consider the input firing rates to be constant over visits (in first approximation). This implies $\langle \nu_j^A \nu_j^B \rangle_j = \langle \nu_j^2 \rangle_j$, which then gives:

$$\rho_t = \frac{\langle c_j^t c_j^{t+1} \rangle - \alpha^2}{\alpha \left(1 - \alpha\right)}, \tag{35}$$

where $t$ stands for the time of the visit. Eq. (35) can be made more explicit by expressing $\langle c_j^t c_j^{t+1}\rangle$ in terms of probabilities. In fact, since $c_j$'s are binary matrix elements, we can write:

$$\langle c_j^t c_j^{t+1}\rangle = \text{Pr}\left(c_j^t = 1, c_j^{t+1} = 1\right) = \text{Pr}\left(c_j^{t+1} = 1 \middle| c_j^t = 1\right)\text{Pr}\left(c_j^t = 1\right). \tag{36}$$

Since we have $\text{Pr}\left(c_j^t = 1\right) = \alpha$, we then have

$$\rho_t = \frac{\text{Pr}\left(c_j^{t+1} = 1 \middle| c_j^t = 1\right) - \alpha}{1-\alpha}. \tag{37}$$

If there is no plasticity, then $\text{Pr}\left(c_j^{t+1} = 1 \middle| c_j^t = 1\right) = 1$ and $\rho_t = 1$, while if there is a complete rewiring $\text{Pr}\left(c_j^{t+1} = 1 \middle| c_j^t = 1\right) = \alpha$ and $\rho_t = 0$.

**Case 2.** In this case, we assume that the connectivity matrix is the same for the different visits, and the input patterns are partially correlated. In the data we have, this would for example be the case of considering the two running directions as different environments. We then have here that $\langle c_j^A c_j^B\rangle = \alpha$, so that the correlation is

$$\rho_{LR} = \frac{\langle \nu_j^L \nu_j^R\rangle_j}{\langle \nu_j^2\rangle_j}, \tag{38}$$

where $L$ and $R$ stands for left and right. Input neurons $j$ are either silent or active with firing rate $\nu$, so that we have $\langle \nu_j^2\rangle_j = f\nu^2$, where $f$ is the sparsity of the input layer (fraction of active cells). As for the previous case, we can write more explicit the correlation using probabilities:

$$\rho_{LR} = \frac{\nu^2 \text{Pr}\left(\nu^L = \nu, \nu^R = \nu\right)}{f\nu^2} = \text{Pr}\left(\nu^L = \nu \middle| \nu^R = \nu\right); \tag{39}$$

the correlation between running directions is just the probability that given an input cell is active in one running idrection, it is active also in the other.

**Case 3**. Since environments are assumed to be orthogonal or independent, then we have $\langle \nu^A \nu^B\rangle = f\nu^2$, so that we get for the correlation

$$\rho_{AB} = \frac{f\left(\langle c^A c^B\rangle - \alpha^2\right)}{\alpha\left(1-\alpha\right)}. \tag{40}$$

We summarize here the three correlations found:

- $\rho_t = \dfrac{\text{Pr}\left(c_j^{t+1} = 1 \middle| c_j^t = 1\right) - \alpha}{1-\alpha}$,
- $\rho_{LR} = \text{Pr}\left(\nu^L = \nu \middle| \nu^R = \nu\right)$,
- $\rho_{AB} = \dfrac{f\left(\langle c^A c^B\rangle - \alpha^2\right)}{\alpha\left(1-\alpha\right)}$.

#### Statistics of connectivity in Hebbian plasticity model

We assume that the learning process has gone on long enough so that the connectivity has reached a stationary state and hence is independent of the initial configuration. Then we consider the encoding of one pattern and calculate the mean and the variance in the number of inputs to a cell in the postsynaptic layer (CA1). We assume $N$ neurons in the presynaptic layer, e.g. EC, and patterns with sparseness $f_{\text{pre}}$ and $f_{\text{post}}$ respectively. The synapse between two co-active cells is potentiated with probability $p_+$, and a synapse between one active and one inactive cell is depressed with probability $p_-$. Synapses between inactive cells are not updated. We consider binary synapses such that the connection from cell $j$ in the presynaptic layer to cell $i$ in the postsynaptic layer is $c_{ij} \in \{0, 1\}$. The degree of a cell $i$ can be written

$d_i = \sum_{j=1}^{N} c_{ij}$. The mean degree in the network is then $\mu_d = \langle d_i \rangle = N\langle c_{ij} \rangle$, and the average over synapses can be written as

$$\langle c_{ij} \rangle = f_{\text{post}} \langle c_{ij} \rangle_{i \in \Omega_f} + (1 - f_{\text{post}}) \langle c_{ij} \rangle_{i \notin \Omega_f}, \tag{41}$$

where $\omega_f$ is the set of neurons which are part of the current, activated pattern. By considering the probability that a synapse is in the potentiated state depending on whether or not the neuron is part of the pattern, one arrives at

$$\langle c_{ij} \rangle = \frac{f_{\text{pre}} f_{\text{post}} p_+}{f_{\text{pre}} f_{\text{post}} p_+ + \big[f_{\text{pre}}(1 - f_{\text{post}}) + f_{\text{post}}(1 - f_{\text{pre}})\big] p_-} \tag{42}$$

The variance in the degree $\sigma_d^2 = \langle d_i^2 \rangle - \mu_d^2$. The first term can be written

$$\langle d_i^2 \rangle = \langle \sum_j c_{ij}^2 + \sum_j \sum_{k \neq j} c_{ij} c_{ik} \rangle = N \langle c_{ij}^2 \rangle + N(N-1) \langle c_{ij} c_{ik} \rangle$$

The first term $\langle c_{ij}^2 \rangle = \langle c_{ij} \rangle$, while the second term can be calculated by considering the probability of both connections being potentiated as a function of whether or not the pre- and postsynaptic cells are active in the current pattern, or not. The result is

$$\langle c_{ij} c_{ik} \rangle = \frac{f_{\text{pre}} f_{\text{post}} p_+ \Big( f_{\text{pre}}(p_+ + 2(1 - p_+)\langle c_{ij} \rangle) + 2(1 - f_{\text{pre}})(1 - p_-)\langle c_{ij} \rangle \Big)}{1 - f_{\text{post}} \big( f_{\text{pre}}(1 - p_+) + (1 - f_{\text{pre}})(1 - p_-) \big)^2 - (1 - f_{\text{post}})(1 - f_{\text{pre}} p_-)^2} \tag{43}$$

Finally, the autocorrelation of the degree can be written

$$\rho_d = \frac{\langle d_i^t d_i^{t-1} \rangle - \mu_d^2}{\sigma_d^2}, \tag{44}$$

where we can write $\langle d_i^t d_i^{t-1} \rangle = N \langle c_{ij}^t c_{ij}^{t-1} \rangle N(N-1) \langle c_{ij}^t c_{ik}^{t-1} \rangle$. The two covariances can, once again, be calculated by considering the probability that the product is equal to one depending on the participation of the cells in the patterns at time $t-1$ and $t$. The result is

$$\langle c_{ij}^t c_{ij}^{t-1} \rangle = \langle c_{ij} \rangle \Big( 1 - p_-(f_{\text{post}}(1 - f_{\text{pre}}) + f_{\text{pre}}(1 - f_{\text{post}}) \Big), \tag{45}$$

$$\begin{aligned} \langle c_{ij}^t c_{ik}^{t-1} \rangle &= \langle c_{ij} \rangle f_{\text{pre}} f_{\text{post}} p_+ \\ &\quad + \langle c_{ij} c_{ik} \rangle \Big( 1 - f_{\text{pre}} f_{\text{post}} p_+ - p_-(f_{\text{post}}(1 - f_{\text{pre}}) + f_{\text{pre}}(1 - f_{\text{post}}) \Big) \end{aligned} \tag{46}$$
